# Unexpected levels of antibiotic resistance in marine bacterial communities not solely explained by urban river discharge

**DOI:** 10.1101/2024.11.15.623850

**Authors:** Brandon Chan, Carolyn Xue, Natalie Lozano-Huntelman, Karina Jimenez, Jennifer Jay, Pamela Yeh, Colin Kremer

**Affiliations:** Department of Ecology and Evolutionary Biology, University of Connecticut, Storrs, CT, USA; Department of Ecology and Evolutionary Biology, University of California Los Angeles, Los Angeles, CA, USA; Civil and Environmental Engineering Department, University of California Los Angeles, Los Angeles, CA, USA; Institute of the Environment and Sustainability, University of California Los Angeles, Los Angeles, CA, USA; Santa Fe Institute, Santa Fe, NM, USA

**Keywords:** Microbial communities, antibiotics, antibiotic resistance, marine bacteria, aquatic environments

## Abstract

Driven by anthropogenic pressures, antibiotic resistance has become more prevalent within urban bacterial communities over recent decades. In general, resistance is expected to spread from highly impacted areas into surrounding environments as the spatial patterns of resistance reflect the balance between the ability of dispersal and horizontal gene transfer to spread resistance, and the effects of dilution or selection driven by other environmental or ecological factors that change along these gradients. However, the strength and scale of this effect is less clear, especially in coastal environments. We investigated antibiotic resistance of bacterial communities along a transect from a freshwater site in the Los Angeles River (surrounded by high-density urban development) out into the San Pedro Basin in the Pacific Ocean. We determined the minimum concentration of twelve different antibiotics needed to inhibit the growth of bacterial communities across eight sites. We also examined community composition and the prevalence of known genes related to antibiotic resistance from anthropogenic sources in each sample using 16S rRNA sequencing and qPCR. In contrast to our expectations, marine bacteria communities displayed higher resistance than the freshwater LA River community. Rather than the presence of known resistance genes or proximity to urban impacts, patterns of resistance were more strongly associated with the higher species diversity found in marine communities. These unexpected results open new opportunities to determine why marine bacterial communities possess high resistance and the repercussions for potentially pathogenic bacteria and their impacts on the health of humans and wildlife.

## Importance

Antibiotic resistance is a growing issue for human and environmental health caused by the expanded use of antibiotics in health and agricultural settings. Yet few studies have investigated resistance in marine bacterial communities, and whether it too is being exacerbated by proximity to human influences. We examined resistance to twelve antibiotics in bacterial communities collected along a gradient from a major, highly impacted riverway in Los Angeles County out into the Pacific Ocean. Unexpectedly, diverse marine bacterial communities displayed consistently higher resistance to a range of antibiotics than the freshwater community. Resistance mechanisms often found in urban environments were not present in marine bacteria, indicating that they may harbor unique mechanisms for antibiotic resistance. Future research should investigate the nature of these mechanisms, why marine communities are so resistant, and what the implications are for the health of human and wildlife populations in coastal regions, at the interface of microbial communities from different environments.

## Introduction

The widespread use and overuse of antibiotics has led to a dramatic increase in antibiotic resistance genes in bacteria (D’Costa et al, 2006; Sommer et al., 2009; Frieri et al., 2017; Knapp et al., 2010). Antibiotic resistance has existed for millions of years as one of the most common defense mechanisms used in conflicts between bacteria, arising through mutation, amplified by selection, and spreading via horizontal gene transfer (HGT; the transfer of genetic material from one bacteria cell to another) (D’Costa et al., 2011; Munita & Arias, 2016). In natural systems, adaptations to external stress, like temperature, can also indirectly confer antibiotic resistance (Cruz-Loya et al., 2019, Kamruzzaman & Iredell, 2019). While the original ecological advantage of antibiotic resistance is associated with competition between bacteria, the exponential increase of antibiotic use by humans has caused an increase in resistant bacteria through intensive selection (Aminov, 2009; Toprak et al., 2012; Livermore, 2009; Allen et al., 2010). Resistance is expected to arise with some frequency as a response to selection in cases where antibiotic treatment is intentionally applied. However, widespread misuse/overuse, improper disposal, unintentional contamination, and lack of metabolization of drugs also introduce antibiotics into environments near urban areas and agriculture, leading to the rise of resistance (Iwu et al., 2020; Halling-Sørensen et al., 1998). Widely used drugs are regularly detected in environments near human populations and even trace concentrations of antibiotic compounds can be enough to select for resistance (Chow et al., 2021; Szekeres et al., 2018; Almakki et al., 2019). Although this has been extensively studied in terrestrial and freshwater systems, very little is known about whether anthropogenic impacts affect antibiotic resistance at the interface of freshwater and marine bacterial communities.

In urban areas, antibiotic pollution, a result of waste from hospitals and industrial antibiotic production, runoff from agriculture, and discharge from wastewater treatment plants, mixes with freshwater environments (Larrson & Flach, 2022; Kraemer et al., 2019; Rizzo et al., 2013; Li et al., 2019). Freshwater environments act as a catchment area for antibiotic pollution, which changes the composition of their bacterial communities and increases the probability for bacteria to acquire resistance from selection or transfer resistance genes to other bacteria through HGT (Aminov, 2009; Amarasiri et al., 2020). This has been shown to increase resistance among bacteria both near the source of pollution and further downstream (Costanzo et al., 2005; Storteboom et al., 2010). Much of this freshwater discharge reaches coastal areas which contain unique community compositions as distinct communities intermix within the surface and are highly subjected to human impacts and environmental factors (Aguiló-Ferretjans et al., 2008; Logares et al., 2009; Nogales et al., 2007; Langenheder & Lindström, 2019; Mobilian et al., 2020). As this discharge containing trace amounts of antibiotics, antibiotic resistance genes, and antibiotic-resistant bacteria reaches the ocean, there is a significant risk of spreading resistance between bacteria from different environments (Hatosy & Martiny, 2015; Nogales et al., 2011; Niu et al., 2016; Zheng et al., 2021). Considering that at least a third of the world’s population lives within 100km of a coast, the influx of these antibiotic pollutants into coastal environments can have serious consequences for marine ecosystems broadly as well as public health, concerning those who use the ocean for commercial and recreational purposes (Brown et al., 2006).

The composition of marine bacterial communities differs from freshwater communities, and may harbor unique functional diversity, including antibiotic resistance phenotypes. Marine bacterial communities often consist of a few classes in the Pseudomonadota (aka Proteobacteria) and Bacteroidota (aka Bacteroidetes) phyla, which include but are not limited to *Alphaproteobacteria*, *Gammaproteobacteria*, and *Flavobacteriia*, and fluctuate in repeated and predictable patterns throughout the water column (Fuhrman et al., 2015; Yeh & Fuhrman, 2022; Paver et al., 2018; Cottrell & Kirchman, 2000; Fuhrman et al., 2006; Hatosy et al., 2013; Cram et al., 2015). In one of the few studies to directly test for antibiotic resistance in marine communities, communities further from shore showed increased resistance phenotypes than near shore (Sizemore & Colwell, 1977). More recently, genotype analysis shows that marine bacteria have acquired a great diversity of putative antibiotic resistance genes, many of which were never previously described in freshwater or terrestrial environments (Hatosy & Martiny, 2015; Cuadrat et al., 2020). While many marine bacteria have different resistance genes than their freshwater and terrestrial counterparts, there is some evidence of the spread of resistance genes from human sources. For example, antibiotic resistance genes and bacteria from human sources have consistently been found in estuaries (Zhu et al., 2017). Human pathogens have also acquired resistance genes from aquatic environments (Finley et al., 2013; Razavi et al., 2017). This is evident in marine bacteria as coastal environments have shown direct evidence of altering the skin and gut microbiome of people (Nielsen et al., 2021; Leonard et al., 2018). However, it remains generally unclear how resistant marine bacterial communities are to antibiotics, and to what extent this is exacerbated by antibiotic pollution and bacterial dispersal from urban areas into coastal environments.

We sought to understand if and how the prevalence of antibiotic resistance in bacterial communities’ changes along a transect from freshwater to marine environments in a densely populated area. Specifically, we sampled microbial communities from the Los Angeles (LA) River, a highly urbanized river surrounded by the second largest city in the United States, out into the Pacific Ocean, following a transect into the San Pedro Channel sampling stations along the San Pedro Ocean Time-series (SPOT). We experimentally tested twelve antibiotics to determine the minimum amount needed to inhibit growth in these bacterial communities. Across this gradient, we also characterized changes in the composition of bacterial communities (based on 16S rRNA sequences) and the prevalence of marker genes for antibiotic resistance and urbanization. We predicted that antibiotic resistance and the corresponding resistance genes would be highest closest to the LA River and decrease further from shore. However, we found that antibiotic resistance increased with distance from shore, often rising abruptly at the boundary between marine and freshwater communities, which was correlated with increased species diversity.

## Methods

### Study Site and Sample Collection

The LA River is a unique and heavily modified waterway that flows through Los Angeles County, the most populous county in the United States (US Census, 2020). Historically, the river was a dynamic floodplain that changed course depending on flooding events. However, 51 miles of the LA River have been channelized since 1960 to mitigate flood damage, with only three sections of the river remaining unpaved today (Gumprecht, 1997). Currently, the LA River serves an important purpose as one of the main waterways in Los Angeles County, especially for wastewater treatment discharge, which comprises up to 70% of its total flow (Wolfand et al., 2022). However, sewage spills into the LA River and the surrounding area are common, increasing the risk of antibiotic-resistant bacteria and resistance genes mixing with the freshwater bacterial communities. In April 2023, 250,000 gallons of untreated sewage spilled into the LA River, prompting beach closures throughout the city of Long Beach (Goldberg, 2023). A much larger spill in December 2021 saw nearly 8.5 million gallons of sewage spill into the Port of Long Beach, causing major beach closures throughout Los Angeles and Orange County. In general, sewage spills can have major economic repercussions in the tens of millions of dollars lost from illness alone (Given et al., 2006). With the recurring events of sewage spills in the area, we selected the LA River and locations directly away from the mouth of the river as our primary sampling locations.

We collected a total of eight 2 L samples of surface water along a transect from the LA River to the San Pedro Basin (*Fig. 1*). Two samples were collected from the LA River at approximately 6 km inland (Site 1) and at the mouth of the river in Long Beach (Site 2) during an outgoing tide on September 14, 2022. We sampled once within the Port of Los Angeles (Site 3) while five additional samples were collected from sites approximately every 5.5 km from the San Pedro Basin in the Pacific Ocean along a 24 km transect aboard the R/V *Yellowfin* as part of the SPOT cruise (Yeh & Fuhrman, 2022). Latitude, longitude, distance from site 1, and time of collection were recorded at each sampling site along with the water properties of salinity, pH, and temperature (*A1*). Based on salinity levels, in all subsequent text and analyses, we refer to Site 1 as a ‘freshwater’ sample and all other sites as ‘saltwater’. We also calculated the linear distance between site 1 and all other sites for statistical analysis in the *geosphere* package (1.5-18) in R (Hijmans, 2022). Of the 2 L of water collected from each of the eight sites, 100 mL were set aside and split into four replicates of 25 mL, creating 32 bacterial community samples. These samples were then used to inoculate with Luria Broth (LB) liquid media made with either pure freshwater or artificial saltwater (34 PSU) as solvent, corresponding to site salinity levels. These communities were cultured overnight (∼12 hours), then each culture was mixed with 50% glycerol to create 8 mL of a 25% glycerol solution, stored in cryotubes, and frozen at -80°C to preserve stocks. The remaining 1.9 L of the collected water samples was split into duplicates, and each 950 mL duplicate was filtered to concentrate bacteria for DNA extraction and sequencing. Samples were filtered through a 10 µm polycarbonate filter (Millipore) to remove larger particles and eukaryotic cells, then filtered again through a 0.22 µm mixed cellulose ester filter (Isopore). Filtrate was discarded, and each of the 16 membranes were placed in 50 mL of MilliQ water to resuspend the particles that remained on the membranes. These filters were then stored and used for DNA extractions.

**Fig. 1.**
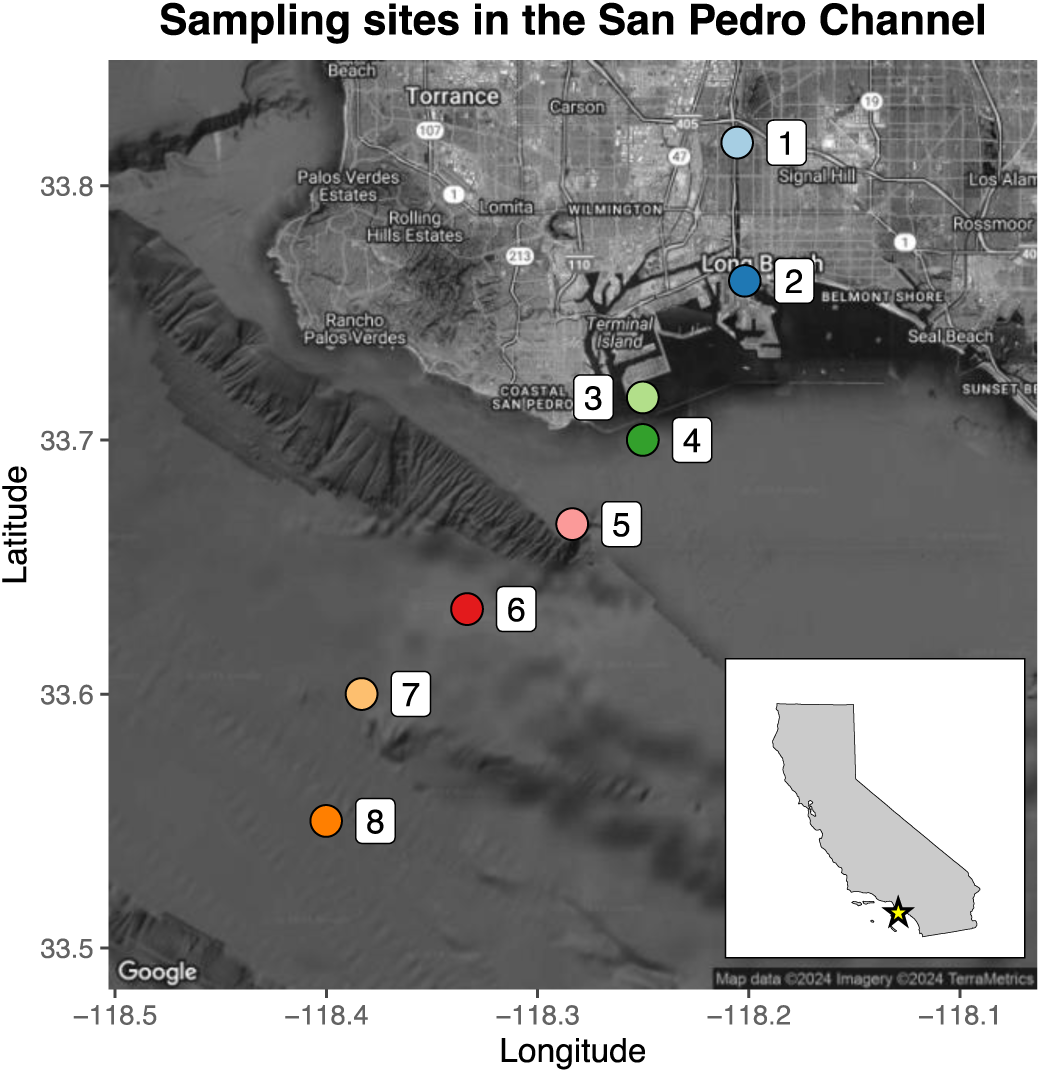
Map of sampling sites located in Long Beach, CA, USA. Sites 1 and 2 were sampled along the Los Angeles River while sites 3-8 were sampled along the September 2022 SPOT cruise.

### Minimum inhibitory concentration (MIC) experiments

The frozen stocks of bacteria from each site, containing subsamples of the bacterial community, were thawed and placed in appropriate LB media (fresh or saltwater) for culturing. Approximately 20 μL of each bacterial sample were placed in 20 mL of LB, incubated for 16-20 hours overnight, then 1:1000 dilutions of these cultures were used for standard experiments to quantify the minimum inhibitory concentration (MIC) for each of twelve antibiotics (*Table 1*) applied to every community. The MIC is the concentration of drug needed to inhibit visible growth by 95% (Kowalska-Krochmal & Dudek-Wicher, 2021). Twelve antibiotics were selected in order to span a variety of drug classes and mechanisms, and for their clinical relevance (*Table 1*). Incubation temperatures of 24°C for sites 3-8 and 30°C for sites 1 and 2 were determined from physical conditions measured at each site. For each site and replicate, a 96-well plate was used to conduct a 20 step two-fold dilution for each antibiotic where the starting concentration was 2000 mg/mL, except for clindamycin and streptomycin which had a higher starting concentration of 16,000 mg/mL. Each plate also had a positive control with cells in LB media with no antibiotic and a blank (a negative control) of only LB media. For each plate, 100 μL of the diluted bacterial cultures were added to every well, except for a column designated as the blank. Plates were allowed to grow overnight (∼12 hours), with optical density at 600 nm measured using a microplate reader the following day to determine growth. The absorbance of a 96-well plate containing only antibiotic solutions, LB media, and no bacteria was also measured to subtract away background absorbance values created by the color of LB media or antibiotic solution. To ensure salinity did not interfere with the effectiveness of the antibiotics, we exposed lab strain *Escherichia coli* in saltwater LB media to a subset of drugs to compare against previous *E. coli* strains grown in freshwater LB media (Bullivant et al., 2024). Overall, MIC values to antibiotics still showed similar patterns for cell growth across different drug types and classes, independent of salinity (*A5*).

**Table 1.** All drugs used in the MIC experiments along with their drug class and mechanism for inhibiting growth.

| Antibiotic | Abbreviation | Class | Mechanism |
| --- | --- | --- | --- |
| Tetracycline | TET | Tetracycline | Inhibition of protein synthesis by targeting the 30S subunit (Grossman, 2016; Forge & Schacht, 2000; Neu, 1976) |
| Gentamicin | GEN | Aminoglycoside |  |
| Streptomycin | STR |  |  |
| Tobramycin | TOB |  |  |
| Clindamycin | CLI | Lincosamide | Inhibition of protein synthesis by targeting the 50S subunit (Spížek & Řezanka, 2017; Vázquez-Laslop & Mankin, 2018) |
| Erythromycin | ERY | Macrolide |  |
| Ampicillin | AMP | Beta-lactam | Inhibition of cell wall synthesis (Oates et al., 1988) |
| Cefoxitin | FOX |  |  |
| Ciprofloxacin | CPR | Fluoroquinolone | Disruption of DNA gyrase function (Blondeau, 2004) |
| Levofloxacin | LVX |  |  |
| Sulfamonomethoxine | SLF | Sulfonamide | Inhibition of folic acid synthesis (Sköld, 2000) |
| Trimethoprim | TMP | Diaminopyrimidine | Inhibition of folic acid metabolism (Fernández-Villa et al., 2019) |

We fit dose response curves to the growth data using the *drc* package in R and MIC values were calculated for 95% inhibition of growth compared to a positive control (Ritz et al., 2015). For comparative analysis, we used MICs set at 95% as the highest level of inhibition detectable by optical density. When exposed to ampicillin, marine bacteria continued to show strong growth even when the drug was applied at its highest soluble concentration. This demonstrated extreme resistance to ampicillin as we were unsuccessful in determining the exact MIC value, so ampicillin was removed from subsequent analyses.

### Thermal performance curve (TPC) experiments

We conducted an experiment to determine whether communities that showed high antibiotic resistance also had higher thermal tolerance. We quantified the thermal performance curve (TPC) of communities, including their optimum temperatures (T_opt_), where growth was maximized, and the temperature range where growth was positive (T_min_ and T_max_). All bacterial culture samples were thawed and placed in their respective LB media (made with freshwater or artificial seawater). Samples were allowed to grow overnight for approximately 16 hours. Approximately 10^5^ cells of each sample were then inoculated into 100 μL of LB media in a single well of a 96-well plate. Each sample was done in triplicate. The resulting plate was used as a template to create the experimental TPC plates via pin transferring the inoculum (1 uL per transfer). A total of 22 TPC plates were created where each individual plate went into an incubator set at a different temperature. Incubators were set with a temperature within the range of 10°C to 52°C, in increments of 2°C. Plate absorbances were read every 2 hours for 24 hours. Growth curves were fitted to absorbance data over time using the *growthTools* package (Kremer, 2020), yielding estimates of exponential growth rate. Thermal performance curves were then fitted using the Norberg model to obtain estimates of T_opt_, T_min_, and T_max_ (Norberg, 2004).

### DNA extraction, sequencing, and qPCR analysis

We extracted DNA from samples using a Qiagen DNEasy Blood & Tissue kit. Two extractions were performed per water sample from each site, making a total of 16 DNA samples. Eluted DNA concentration was quantified using a Qubit fluorometer, and concentrations per sample ranged from 0.04 to 0.5 µg/µL.

To characterize bacterial community composition, one set of the replicate DNA samples were shipped to the Microbial Analysis, Resources, and Services facility (University of Connecticut) for 16S rRNA Amplicon sequencing. Library preparation, alignment, and 3% OTU classification were performed using the *mothur* pipeline (Schloss et al., 2009). Community composition at the phyla level was visualized using ‘TAXONOMY’ format files produced by the pipeline and visualized using R. OTU tables produced by the pipeline were further analyzed and visualized using the *vegan* package in R (Oksanen et al., 2024). Taxa defined as unique OTUs were used to determine Shannon diversity Index and Jaccard dissimilarity Index for alpha and beta diversity, respectively. Indicator species, found as OTUs with the highest association to a group, were found using the *indicspecies* package in R (De Cáceres & Legendre, 2009).

To determine the prevalence of focal antibiotic and marker genes, the second set of DNA samples was used for qPCR analyses. The antibiotic resistance genes that were amplified were *tetA*, *intI1*, *sul1*, and *blaCTX-M* (*Table 2*). Target genes were chosen based on their known abundance around anthropogenic areas and their clinical relevance for qPCR analysis from urban wastewater samples (Pruden et al., 2013, Echeverria-Palencia et al., 2017). The 16S rRNA gene was also amplified to quantify total bacterial abundance, which was used to normalize antibiotic resistance gene abundance. All assays were performed in MicroAmp Fast 96-Well Reaction Plates (Applied Biosystems), with template DNA loaded in triplicates, and all plates were run using the instrument StepOne Plus Real-Time PCR System (Applied Biosystems). Target gene sequences reported from the WRF 5052 were used to synthesize DNA fragments (IDT technologies) to serve as positive controls. These fragments were serially diluted to create a 7-point standard curve, from a starting concentration of 10^6^ or 10^5^ ng/mL to 1 or 1/10 ng/mL. Further details on qPCR gene targets, primers, and reaction conditions can be found in supplemental information (*S13-14*). One gene, *blaCTX-M*, failed to amplify and was therefore excluded from the results.

**Table 2.**
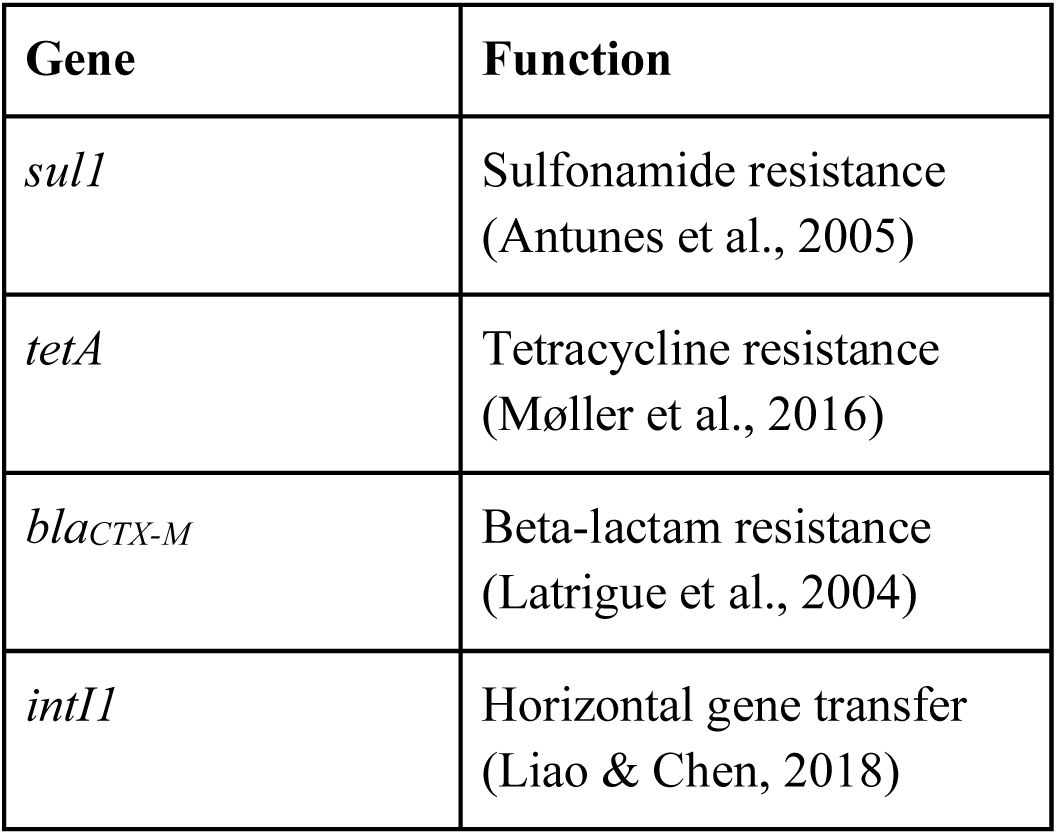
Target genes for qPCR and their gene function.

### Statistical Analyses

#### General MIC

To determine whether some communities tended to be more resistant overall than others to antibiotics across the full range of eleven drugs we considered (excluding ampicillin, see above), we created a general MIC metric. This metric (equivalent to a Euclidean distance) is defined as:

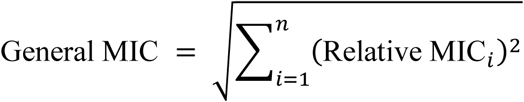

where *i* is the index for each of the *n* = 11 drugs, and relative MIC (ranging from 0 to 1) is determined by dividing the community’s MIC value for a particular drug by the maximum MIC value observed for that drug across all communities. Communities with no resistance to any of the focal drugs would have a general MIC value near zero, whereas communities with maximal resistance to all of the drugs considered would have a general MIC value of 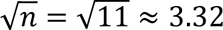. Note, however, that this metric does not discriminate well between situations where a community may have high resistance to a few specific drugs or moderate to low resistance to many drugs.

We used a Monte Carlo approach to calculate uncertainty for the general MIC metric, arising from underlying uncertainty in our MIC estimates for each drug. Using the estimated mean and standard deviation from each replicate, drug, and site, we simulated new MIC values from a normal distribution and then calculated a new general MIC value. We repeated this process 10,000 times. From these values, we then calculated 95% confidence intervals (as the 2.5^th^ and 97.5^th^ quantiles).

Next, we analyzed variation in the mean values using a generalized additive model (GAM) to compare the effect of distance from site 1 on general MIC values. To facilitate interpretation of the statistical analysis, we incorporated distance effects as both a parametric, linear term and a higher order, non-parametric, thin-plate smoothing term. To avoid redundancy, we omitted the linear basis function of the thin-plate smoother. To avoid overfitting, given the small number of observations, we used restricted maximum likelihood estimation and set the initial number of knots to five. We fit All GAM models using the *mgcv* package and visualized using the *ggplot2* package (Wood, 2011; Wickham, 2016).

#### Drug-specific statistical analyses

For each drug, we tested whether MIC values differed with distance and species diversity using a GAM. Prior to fitting the model, we log10-transformed all MIC values to conform to assumptions of normality. As described in the previous section, we used a parametric linear term and non-parametric, thin plate smoothing term. Finally, to compare MIC values to qPCR results, we fit a linear regression to the data.

All summaries of model analyses can be found in the appendix (*A6-12*). We used R to perform all statistical analyses and associated code and data are provided (V4.4.0, R Core Team, 2024).

## Results

### Changes in MICs

Regarding the overall resistance of bacterial communities (as measured by general MIC across all drugs), we found no significant effect of distance along our transect from the LA River into the San Pedro basin (*Fig. 2A*). However, for ten specific drugs, we found significant spatial structure driven by a mixture of linear and nonlinear trends (streptomycin (STR) was the only exception). For six drugs, we found both a positive linear trend with distance and a significant nonlinear trend (cefoxitin (FOX), gentamicin (GEN), erythromycin (ERY), levofloxacin (LVX), tetracycline (TET), and tobramycin (TOB)). Of the remaining drugs, three displayed only a significant nonlinear trend (clindamycin (CLI), ciprofloxacin (CPR), trimethoprim (TMP)), all of which showed apparently unimodal relationships between MIC and distance. Only one drug, sulfamonomethoxine (SLF), showed an overall negative linear trend with distance from shore (*Fig. 2B*), as well as a significant nonlinear term. For 10 out of 11 antibiotics, the relationship between distance and MIC values is non-linear, and often strongly differentiated between sites 1 and 2. In subsequent sections, we consider whether other properties of these bacterial communities may help explain the patterns displayed by the unexpectedly high MIC values in marine sites. These include whether marine bacterial communities have a higher abundance of known resistance genes from anthropogenic sources (for tetracycline and sulfamonomethoxine), or generally harbor higher bacterial diversity, which might increase the chances that a bacterial strain with resistance is found within a community by chance.

**Fig. 2.**
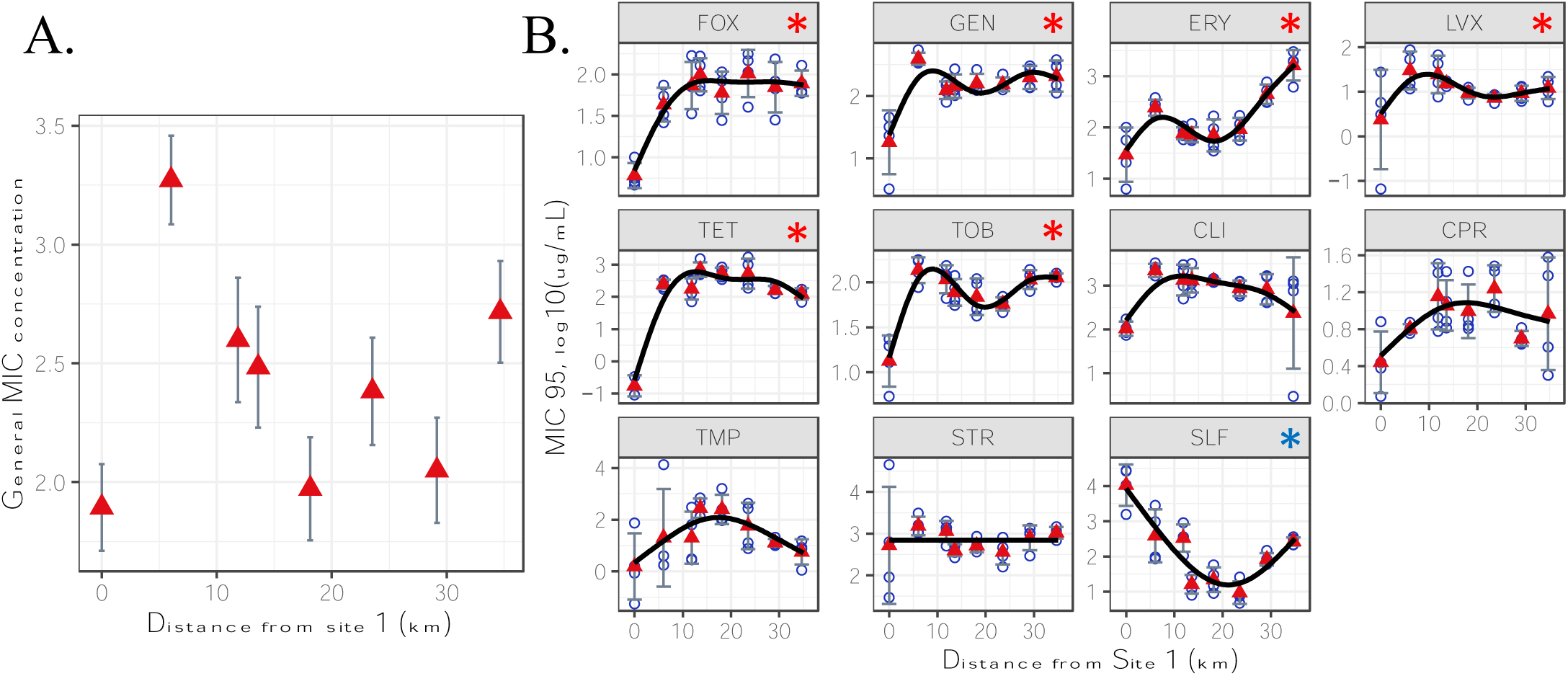
Antibiotic resistance (MIC 95) varies with distance from highly-impacted freshwater site 1 out nto the San Pedro basin. (A) Our general MIC metric across drugs shows no significant trend with distance. (B) However, antibiotic resistance increases with distance for half of the individual drugs including a significant, positive linear trend; red asterisks). Only a single drug (SLF) shows a negative elationship (blue asterisk). Red triangles represent mean MIC values across replicates while the error ars indicate the mean value +/-the standard deviation. Gray bands around trend lines represent 95% onfidence bands. Drug abbreviations can be found in *Table 1*.

### MICs vary with species diversity

Although distance did not structure the overall general MIC values, this metric showed a significantly positive response when compared against bacterial community diversity. Using the general MIC values, alpha diversity had a significant effect of distance on antibiotic resistance (p-value = 0.016) (*Fig. 3A*). More specifically, we found significant spatial structure driven by a mixture of linear and nonlinear trends in nine out of the eleven drugs: FOX, GEN, ERY, LVX, TET, TOB, CLI, CPR, and SLF (*Fig. 3B*). Of these nine drugs, five showed significant structure for both the positive linear trend and the significant nonlinear trend: FOX, GEN, ERY, TET, and TOB. LVX and CLI only showed a significant positive linear trend while CPR and SLF showed a significant nonlinear trend. Only one drug, SLF, showed a negative trend between SDI and MIC values.

**Fig. 3.**
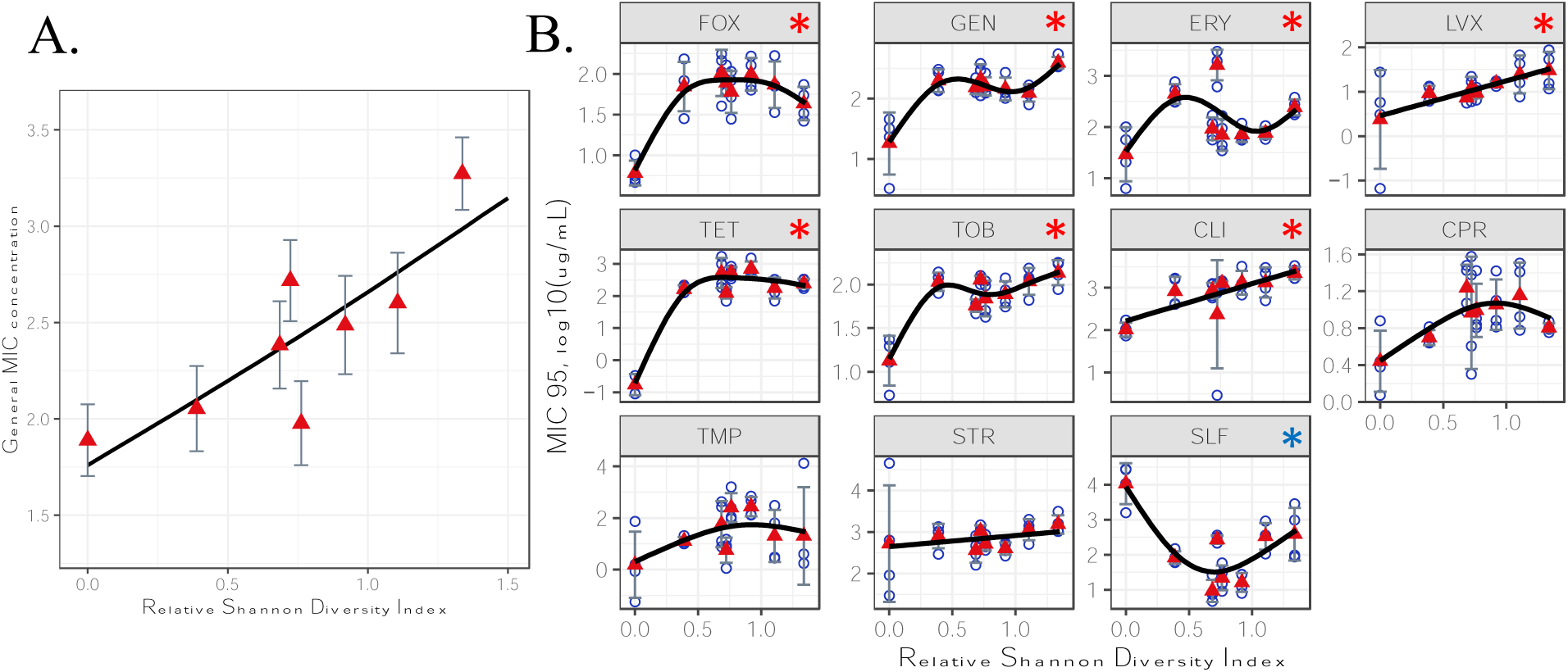
Antibiotic resistance (MIC 95) varies with species diversity (relative Shannon Diversity Index; the observed SDI at each site minus the lowest SDI across sites). (A) Generally, resistance has a significant positive linear relationship with species diversity (p-value = 0.016). (B) Individually, seven drugs show a significant positive linear relationship between species diversity and resistance (red asterisks) while one drug showed a negative relationship (blue asterisks). Red triangles represent mean values while the error bars indicate the mean value +/-the standard deviation. Gray bands around trend lines represent 95% confidence bands. Drug abbreviations can be found in Table 1.

### qPCR results

Sites 2 and 3 had the highest amounts of 16S rRNA, while site 1 had slightly greater abundance compared to sites 4 through 8, indicating the presence of more bacteria (*A3*). The 16S rRNA values were then used to normalize the relative abundance of the three other genes that relate to antibiotic resistance. The *tetA*, *intI1*, and *sul1* genes all showed similar distributions with high relative abundance at sites 1 and 2 and low or undetectable values at the remaining sites (*Fig. 4*).

**Fig. 4.**
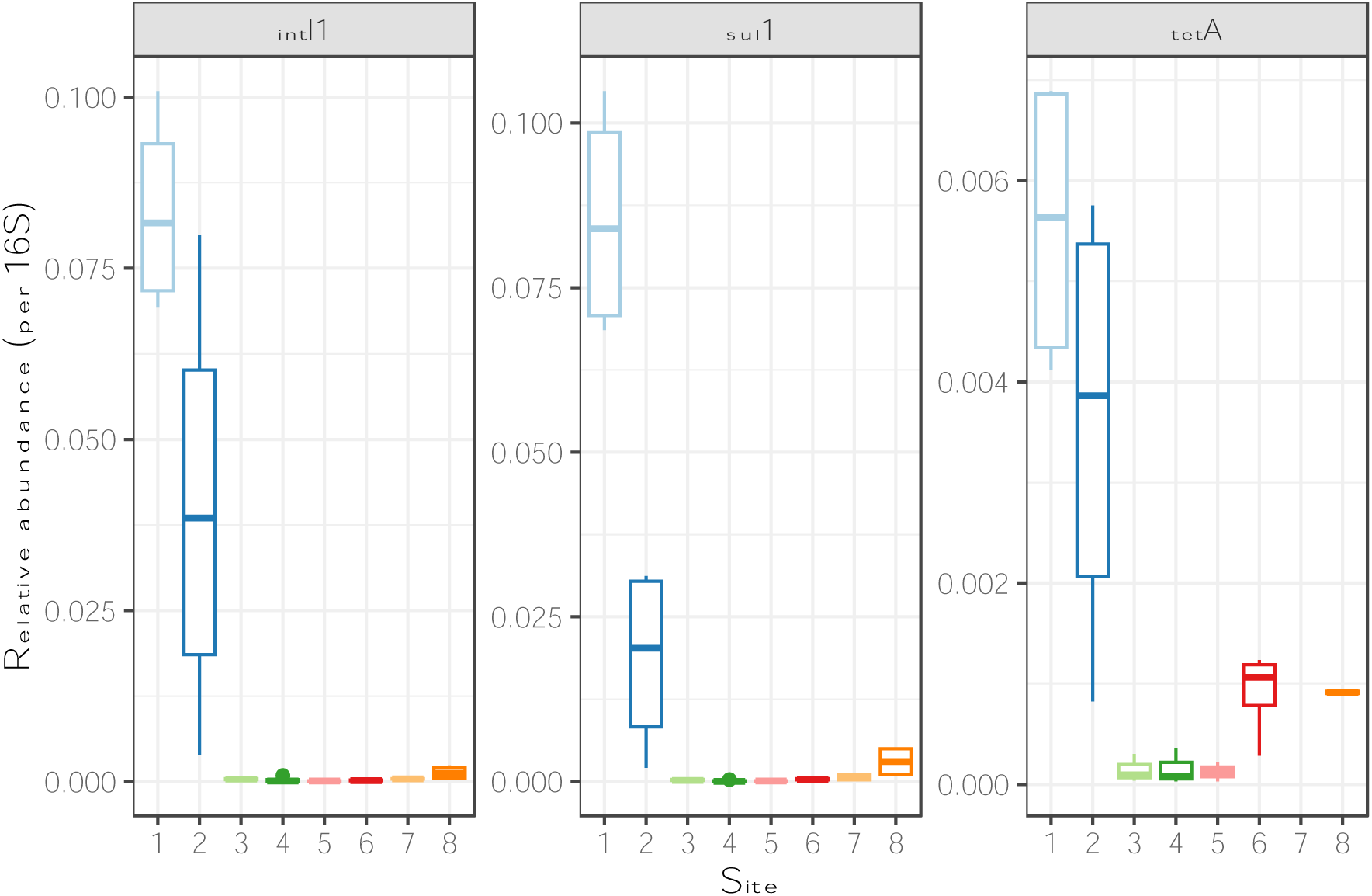
All three genes showed a decreasing pattern of relative gene abundance with distance from shore, with highest abundance closest to shore and little to no abundance within marine communities. qPCR data for three important genes that contribute to horizontal gene transfer (intI1), sulfonamide resistance (sul1), and tetracycline resistance

### Community composition

Bacterial community composition at the level of phyla changed most dramatically between sites 1 and 2, with site 1 being predominantly composed of the phylum Bacteroidota (74.67%) and Pseudomonadota (20.00%) with some bacteria belonging to Actinomycetota (aka Actinobacteriota) (2.31%). In site 2, Pseudomonadota (71.55%) replaces much of the Bacteroidota (21.10%) phyla which is consistent as distance from shore increases. Further from shore, Cyanobacteriota (17-27%) increase in abundance but Pseudomonadota (54.4-61.4%) remain dominant (*Fig. 5*). Alpha diversity at the species level based on unique OTUs, calculated using the Shannon Diversity Index, showed highest diversity in site 2, the saline interface between marine and freshwater at the mouth of the LA River, and lowest diversity upriver in site 1 *(Fig. 6A)*. When examining the overall differences in composition between sites, site 1 differs the most from the other sites, while the sites further out from shore are more likely to cluster together (*Fig. 6B*). Site 2 also differs considerably from both site 1 and the other marine sites, most likely due to the meeting of both freshwater, saltwater, and brackish bacterial species.

**Fig. 5.**
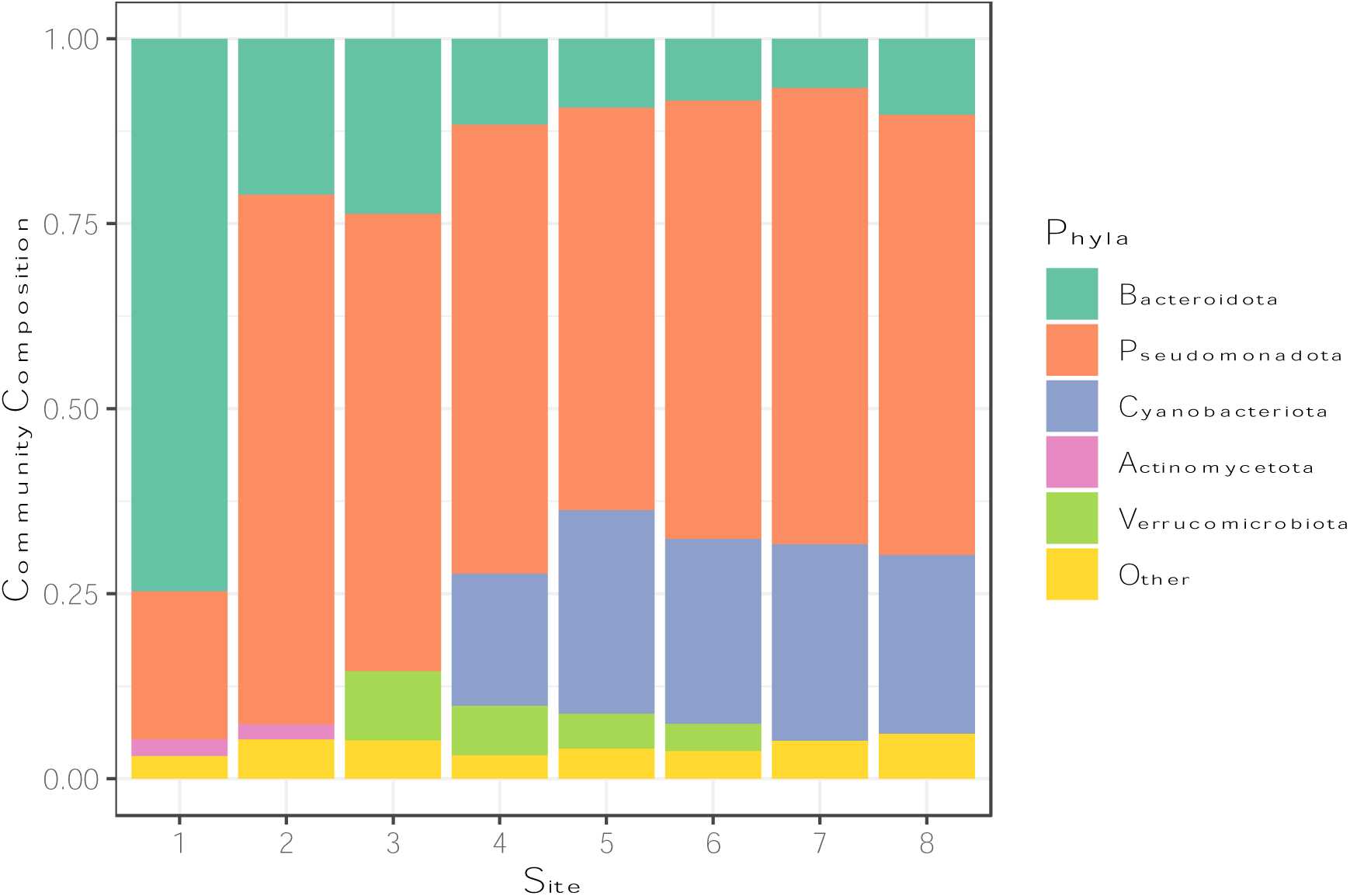
Community composition changes drastically between sites 1 and 2 primarily driven by salinity, while a more gradual change occurs further out from shore with more cyanobacteria becoming present in open ocean environments

**Fig. 6.**
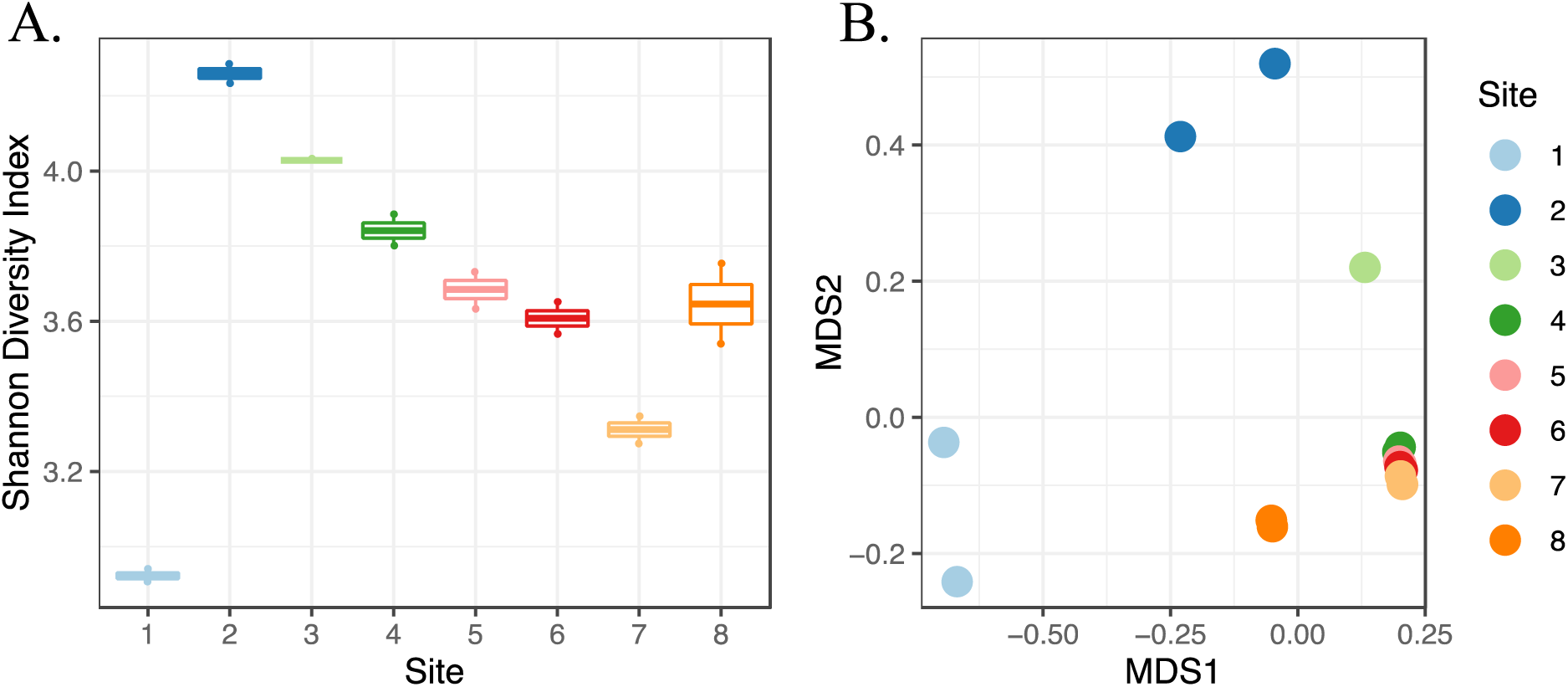
Sites further inland showed larger differences in community composition than sites further from shore. (A) Alpha diversity, as measured by Shannon Diversity Index, was greatest in site 2 while lowest in site 1, with a decreasing trend as distance from shore increased. (B) For specific species, Jaccard dissimilarity showed that community composition becomes more similar further from shore, with sites 1 and 2 being the most different from other sites.

### qPCR and MICs

For qPCR results, we found that the gene abundance of *sul1* displayed a significantly positive correlation with resistance to sulfamonomethoxine (p-value < 0.001) (*Fig. 7*); notably, this was the only drug that showed MIC patterns consistent with our initial expectations regarding anthropogenic influence (*Fig. 4*). However, the gene abundance of *tetA* showed a significantly negative correlation with resistance to tetracycline (p-value = 0.003). Both trends are strongly driven by data from site 1.

**Fig. 7.**
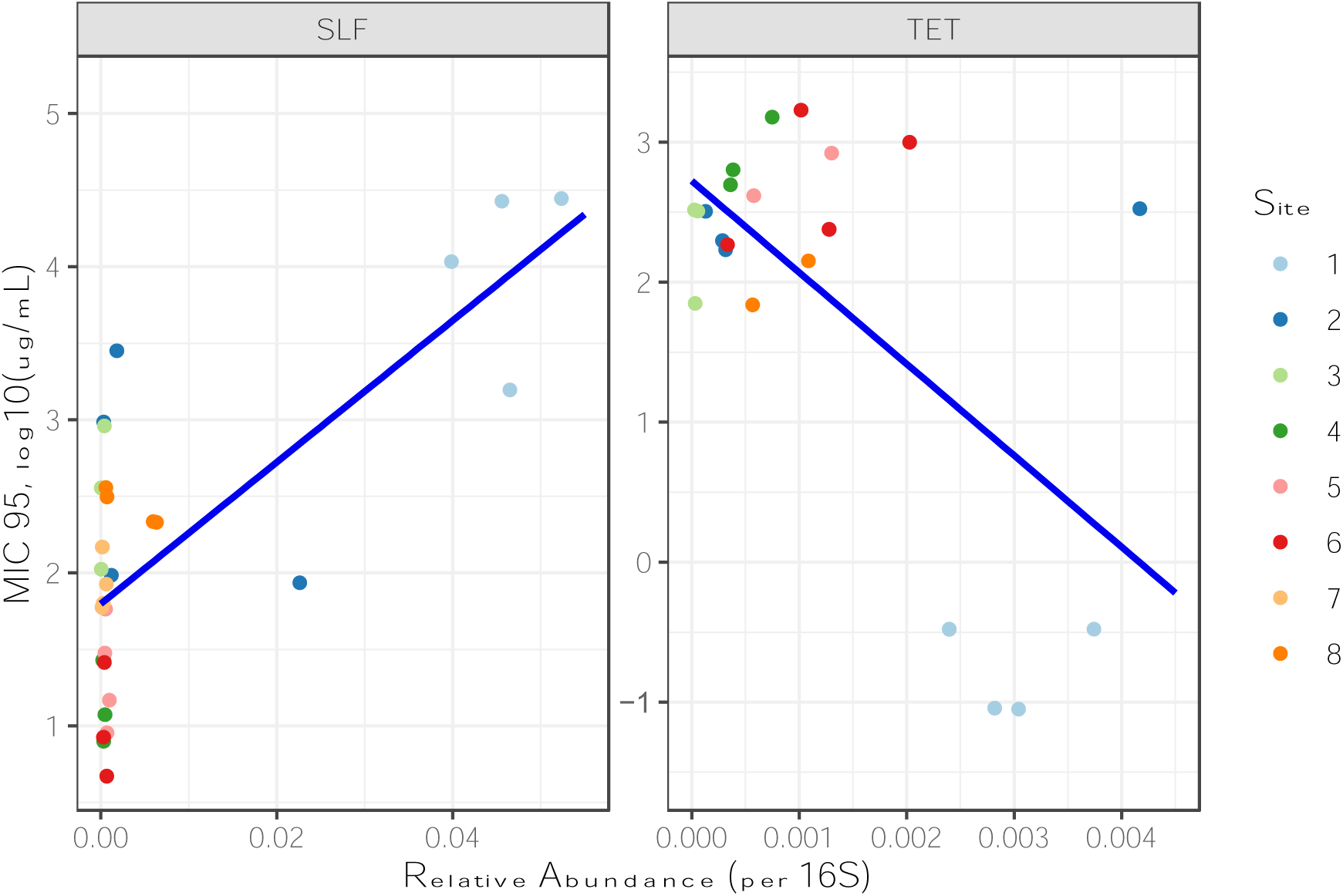
There is a significant linear relationship between gene abundance of sul1 and SLF MIC values (R^2^ = 0.51, p-value < 0.001) and tetA and TET MIC values (R^2^ = 0.34, p-value = 0.003). However, only sul1 corresponds with increased resistance shown in MIC while tetA shows the opposite trend, indicating that resistance is not correlated with gene abundance as expected.

Finally, we also examined the effect of thermal tolerance as a possible mechanism that confers antibiotic resistance (*A2*). We found that there was no significant difference for the thermal maximum between the different sites. While there were minor differences in thermal optimum for site 7 (p-value = 0.02), and the thermal minimum of site 3 (p-value = 0.01) against other sites (*A10-13*), the lack of observable patterns between the relationship of temperature and antibiotic resistance indicates that these effects do not contribute to resistance observed in our samples.

## Discussion

Counter to our expectation that bacteria from sites with more significant anthropogenic impacts (in or near the LA River) should exhibit higher antibiotic resistance, our results show that resistance was significantly greater in marine environments. This held across most — although not all — different antibiotic drugs and drug classes investigated, including drugs with different operating mechanisms (*Fig. 2, Table 1*). This pattern contrasts with our qPCR results, which showed the expected signature of anthropogenic impacts (occurring most prevalently in sites 1 and 2). Indeed, only a single drug (SLF) out of twelve investigated showed the predicted pattern of resistance, which did correlate with qPCR data on the occurrence of an associated known resistance gene. Why are marine bacterial communities more resistant to several different antibiotics than their freshwater counterparts? Although unexpected, this finding is consistent with one historical study (Sizemore & Colwell, 1977). There are several potential explanations, including higher species diversity, adaptation to other environmental factors, and different resistance mechanisms or rates of HGT in marine versus freshwater communities.

For many communities, increased species diversity can often stabilize communities against stressful conditions, especially when those communities are already under some type of stress (Mulder et al., 2001; Jernelöv & Rosenberg, 1976). Microbial communities are no exception as higher species diversity has been shown as a mechanism of bet-hedging against antibiotics (Vega & Gore, 2014). Intuitively, if the broader, regional species pool contains resistant species, then a more diverse site populated by random draws from the species pool would have a greater chance of containing a species resistant to a particular drug. However, microbial communities with lower diversity have also shown an increase in resistance genes (Chen et al., 2019). Our results suggest generally that diversity positively affect resistance, with only a few drugs showing neutral or negative correlations (*Fig. 3*). However, we lack the data to demonstrate which members of the microbial community were responsible for the resistance observed. Given how much community composition differs between freshwater and saltwater environments, identifying whether resistance is shared between many species of bacteria or restricted to only a few species will be an important future step in determining the clinical significance of these resistant bacteria. A broad understanding of species diversity may provide further insights into how strongly antibiotics affect bacterial communities.

Antibiotic resistance may be higher in marine communities if marine bacteria possess unique adaptations to some other environmental stressor, and if these adaptations also confer some level of antibiotic resistance (as is known to occur, Rodriguez-Verdugo et al., 2020). In particular, thermal adaptation has been shown to confer resistance in bacteria to some antibiotics without previous exposure to that drug (Rodriguez-Verdugo et al., 2013; Cruz-Loya, M. et al., 2019). However, we found a lack of unilateral differences in thermal tolerance between the freshwater and saltwater environments. This suggests that antibiotic resistance in marine communities is not explained by marine communities having higher thermal tolerance (*A2*). Indeed, we might expect the freshwater, LA River communities to regularly experience warmer, more variable water temperatures than the marine communities. Some caveats remain: while previous studies have only focused on model species, a community may mask the effects of an individual species that could have a conferred resistance when exposed to antibiotics. Another major environmental condition that differed between our sites is salinity. Salinity is a major factor driving differences in community composition between a river and ocean gradient (Fortunato et al., 2012). The majority of studies on salinity focus on soil bacteria and salt tolerance, finding that highly saline environments contain higher levels of antibiotic resistance genes (Sepúlveda-Correa et al., 2021; Xu et al., 2022; Zhang et al., 2019). Based on our results, it is plausible that cellular mechanisms underlying salt tolerance may also provide antibiotic resistance in bacteria. Saline environments might also drive the evolution of novel resistance mechanisms. This is an exciting avenue for future research.

Marine bacteria have been known to develop novel resistance mechanisms compared to terrestrial bacteria that are constantly exposed to human pollutants (Hatosy & Martiny, 2015). Our qPCR results targeting genes known to associate with anthropogenic impacts mostly align with those findings. All genes that were successfully amplified were disproportionally more common in our freshwater and transitional sites (1 and 2, respectively). For *tetA*, a gene involved in tetracycline resistance, the significant negative linear trend between the abundance of genes and MIC values indicate that another gene or factor must be responsible for the observed resistance in marine bacterial communities. One antibiotic did follow our expectations: freshwater bacterial communities were significantly more resistant to sulfamonomethoxine than marine communities. The abundance of the *sul1* gene, which corresponds to the resistance of the sulfonamide class of antibiotics, follows the phenotypic resistance observed in the MIC experiments. Notably, sulfonamides were the first antibiotics used systematically in clinical and agricultural settings, and as a result, there is widespread resistance (Sköld, 2000, Hutchings et al., 2019). Sulfonamides also are a part of a different class and operating mechanism from the others we considered, and it is possible other drugs of the same class might exhibit similar responses.

Another gene we examined, *intI1*, is involved in mediating horizontal gene transfer and showed high abundance in site 1 with low to no abundance further from shore. Due to its close association with transferring antibiotic resistance in diverse bacterial species, *intI1* is frequently used as an indicator of anthropogenic pollution (Gillings et al., 2014; Ju et al., 2019). However, high resistance profiles in marine sites despite low-to-absent levels of *intI1* may suggest different mechanisms of transmission occurring in marine communities than in freshwater communities. Marine bacteria may be particularly adept at spreading and exchanging antibiotic resistance genes. These bacteria can create gene transfer agents, a HGT mechanism where the bacteria spontaneously produce virus-like particles with their own DNA and can transmit antibiotic resistance at a rate of several hundred times faster than spontaneous mutation of antibiotic resistance genes (McDaniel et al., 2012; Aminov, 2011; McDaniel et al., 2010). These unique mechanisms for gene transfer may play a role in the high resistance observed in marine bacterial communities.

Alternatively, although we think it is unlikely, it is possible that resistance in marine communities is driven by anthropogenic impacts as we initially hypothesized, but that our study design limits our ability to detect this effect. We only have data from a single LA River freshwater site, sampled at a single time point, and additional sampling of the LA River should be a priority for future work. Conditions such as discharge rates, weather, and time of day could all lead to a highly dynamic bacterial community. In contrast to the potentially variable properties of the LA River, the San Pedro basin is much more stable (although far larger), and it has been receiving pollution from LA for many decades, including dichlorodiphenyltrichloroethane (DDT) (Kivenson et al., 2019). The resistance observed in marine communities could be due to this long-term pressure from pollution in general (Baya et al., 1986; Fresia et al., 2019). Additional and repeated sampling across freshwater to marine gradients in urban and less urbanized regions should also be undertaken to understand the generality of our results.

Antibiotic resistance mechanisms present in marine bacteria may have important implications for marine ecosystems and public health, especially for coastal communities. With antibiotics entering the ocean either through pollution or agricultural runoff, many important marine wildlife, from birds to marine mammals are at increased risk of harboring resistant bacteria (Houssain et al., 2022; Rose et al., 2009). The prevalence of resistant bacteria in marine wildlife can act as indicators for the health of the marine ecosystem and the likelihood of acquiring resistant bacteria in nearby human populations (Gross et al., 2022). Although the risk of acquiring a resistant infection from environmental bacteria is unclear in terrestrial environments (Larsson & Flach, 2022; Manaia, 2011), coastal environments have shown more direct evidence of altering human microbiomes (Nielsen et al., 2021; Leonard et al., 2018). Swimming in the ocean was found to be a risk factor for urinary tract infections caused by *E. coli* (Søraas et al., 2013). Sand on coastal beaches also showed to be a direct risk factor for transmission of bacterial diseases like gastrointestinal illness either direct exposure of bacteria living on the sand particles or as a vector for aquatic bacteria transmission (Solo-Gabriele et al., 2016). Notably, site 2 near the mouth of the LA River had the highest bacterial diversity and general MIC values which might be most reflective of beach-adjacent habitats. Therefore, understanding how marine bacteria might interact with other potentially pathogenic bacteria is important for the future public health of coastal communities.

Overall, diverse marine bacterial communities appear surprisingly more resistant to antibiotics than expected, even in comparison to a heavily impacted freshwater bacterial community. The mechanisms underlying this resistance appear to differ from previously observed genes from terrestrial sources, and to be present in different species, as the community composition of freshwater and marine communities varies widely. These unexpected results suggest additional important research, including understanding how resistance arises in marine bacteria and determining the importance of antibiotics to natural microbial interactions within marine environments. Identifying the mechanisms and genes responsible for resistance in marine bacteria is also a priority, especially if these resistance genes can be easily transferred via HGT to potentially pathogenic bacteria, or perhaps into freshwater environments. With the exponential rise in antibiotic resistance exhibited in terrestrial and pathogenic bacteria over time, it is becoming more important to understand how marine bacterial communities influence the resistance of other bacteria along coastal communities.

## Data Availability

All data sets and code have been deposited in a Zenodo repository here: https://doi.org/10.5281/zenodo.14151703 with access to the restricted files here.

## Acknowledgements

We would like to thank the Southern California Marine Institute and Dr. Troy Gunderson for allowing us to join the September 2022 SPOT cruise, the UConn Microbial Analysis, Resources, and Services facility and Dr. Kendra Maas for assistance in DNA sequencing and guidance in analysis, and Hannah Larson and Iris Yang for assisting with field work, sample collection, and experimental work. This material is based upon work supported by the University of California, Los Angeles, the UCLA Institute of the Environment and Sustainability LaKretz student grants, the University of Connecticut, and the National Science Foundation Graduate Research Fellowship under Grant No. (DHE 2136520).

## Disclaimer

Any opinion, findings, and conclusions or recommendations expressed in this material are those of the authors and do not necessarily reflect the views of the National Science Foundation.

## Appendix

**A1.** Table for Sampling site water properties and location collected on September 14, 2022.

| Site | Latitude | Longitude | Distance<br>from Site 1<br>(km) | Temperature<br>(C°) | Salinity<br>(PSU) | Time |
| --- | --- | --- | --- | --- | --- | --- |
| 1 | 33.8168724N | -118.20554991W | 0 | 30 | 0 | 15:08 |
| 2 | 33.7625591N | -118.2019100W | 6.034 | 26.8 | 31 | 14:35 |
| 3 | 33.7167261N | -118.25017889W | 11.852 | 24.4 | 35 | 14:00 |
| 4 | 33.7000496N | -118.25024306W | 13.603 | 24.7 | 35.5 | 13:48 |
| 5 | 33.6669367N | -118.28355333W | 18.133 | 24.3 | 35 | 13:33 |
| 6 | 33.63347833N | -118.33341306W | 23.542 | 24 | 36 | 13:11 |
| 7 | 33.60005333N | -118.38333972W | 29.154 | 24.5 | 35.5 | 12:48 |
| 8 | 33.55002778N | -118.40007944W | 34.661 | 24.5 | 36 | 11:25 |

**A2.**
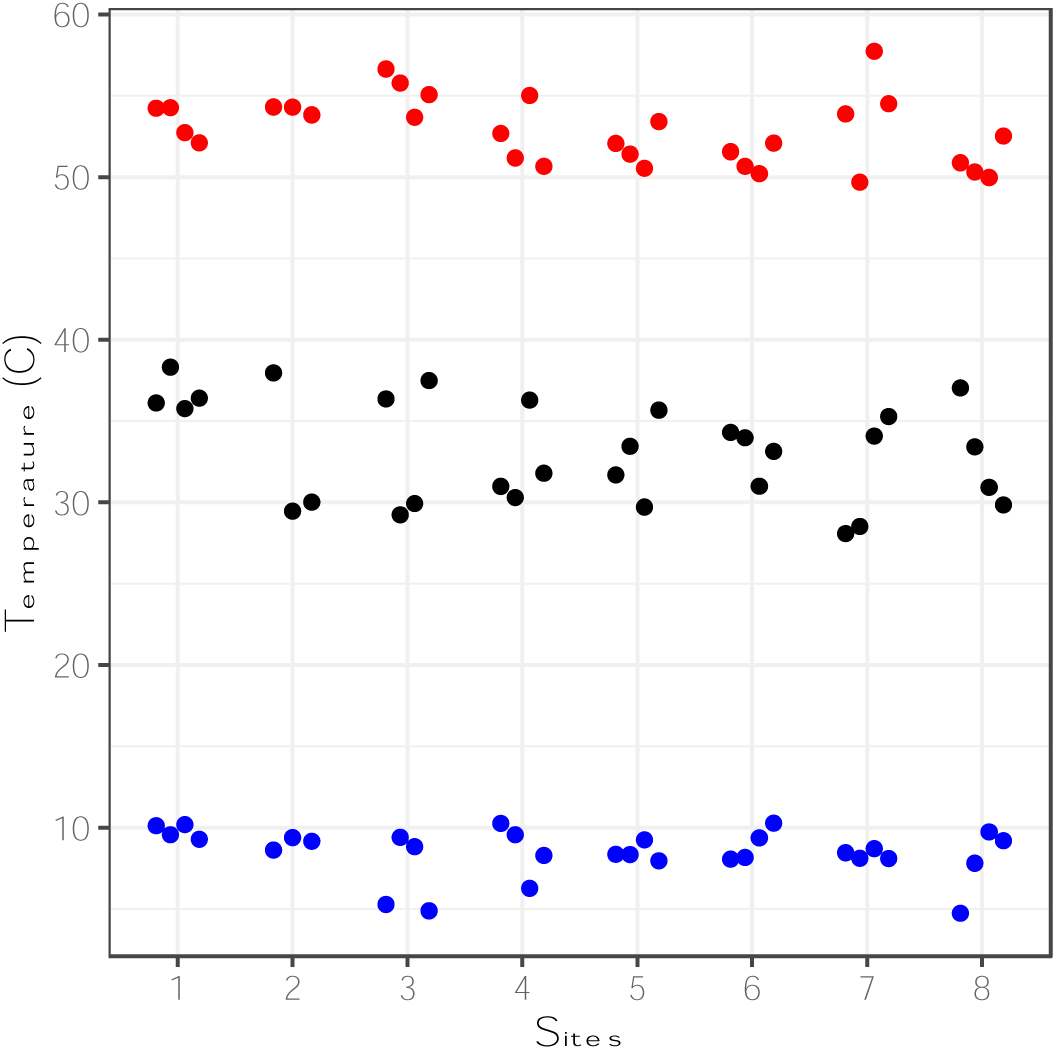
The thermal range of the microbial communities did not significantly differ between sites. Tmax (red), Topt (black), and Tmin (blue) varied slightly between sites but without any discernable pattern to indicate marine bacteria having a higher thermal tolerance.

**A3.**
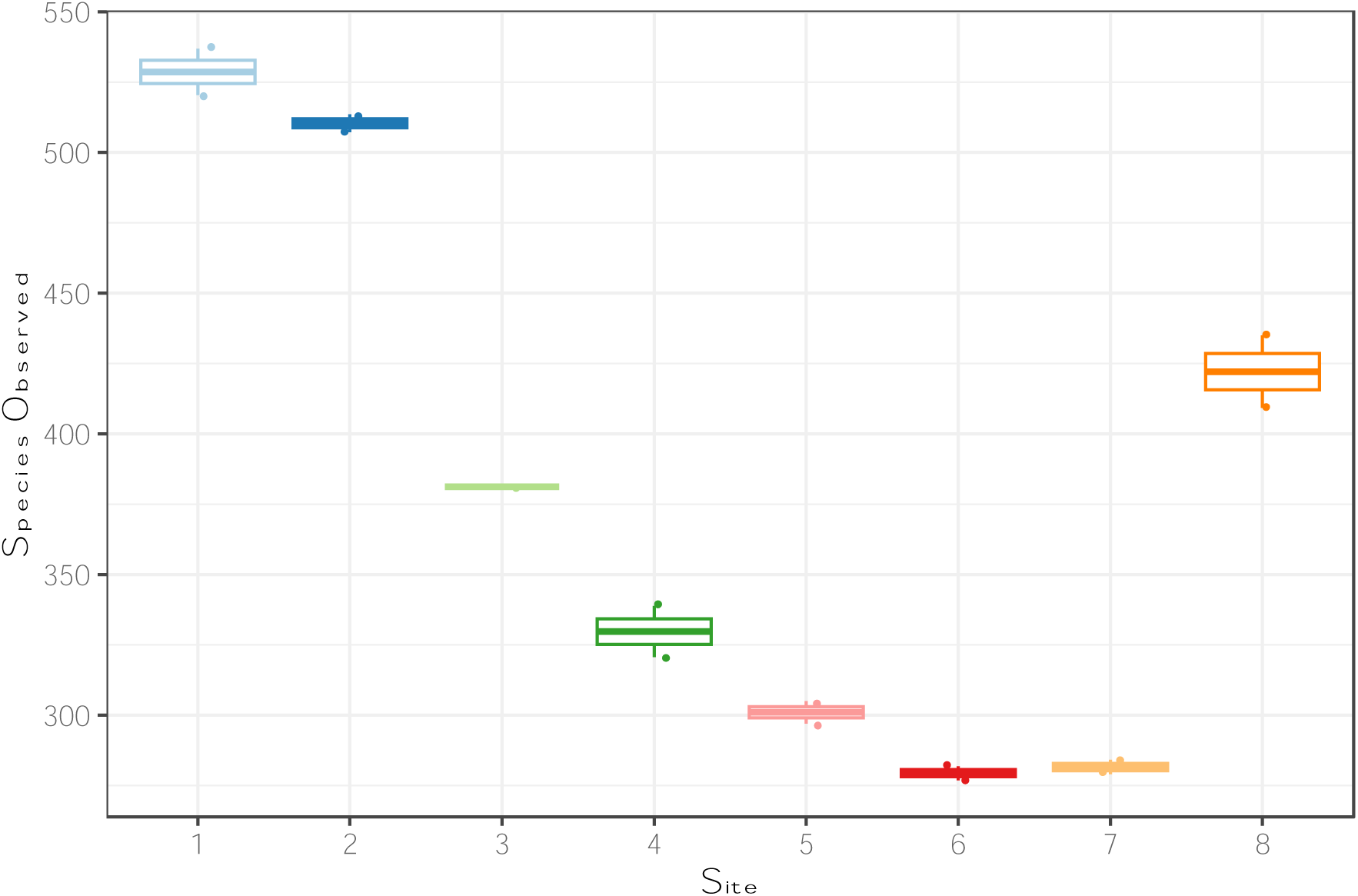
Site 1 has the highest number of OTUs with steep decrease further from shore. However, site 1 also has the lowest species diversity in general

**A4.**
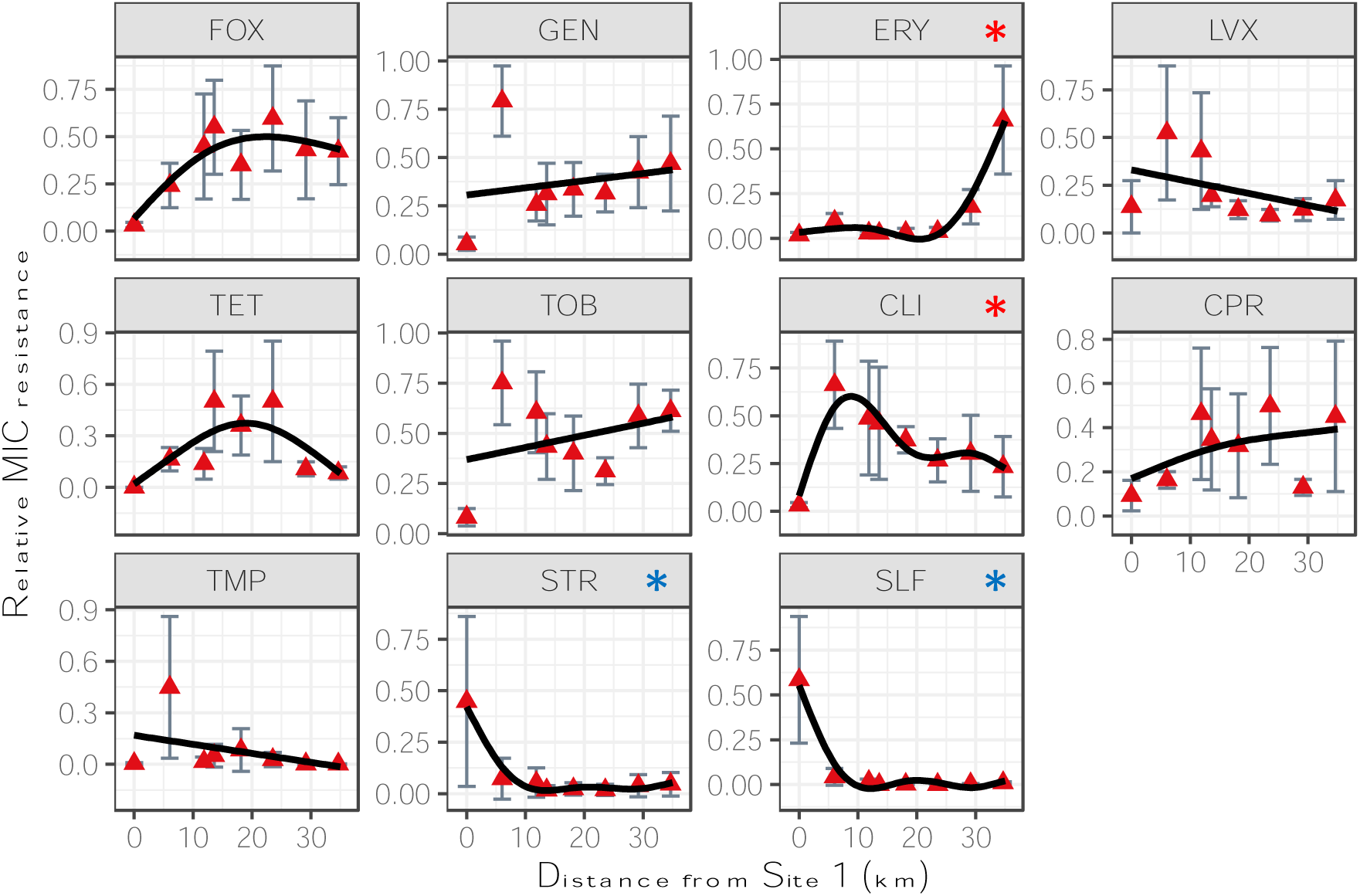
While the overall results show that distance does not significantly contribute to relative MIC resistance, some drugs show a significantly positive relationship (red asterisks while other drugs show a significantly negative relationship (blue asterisks)

**A5.**
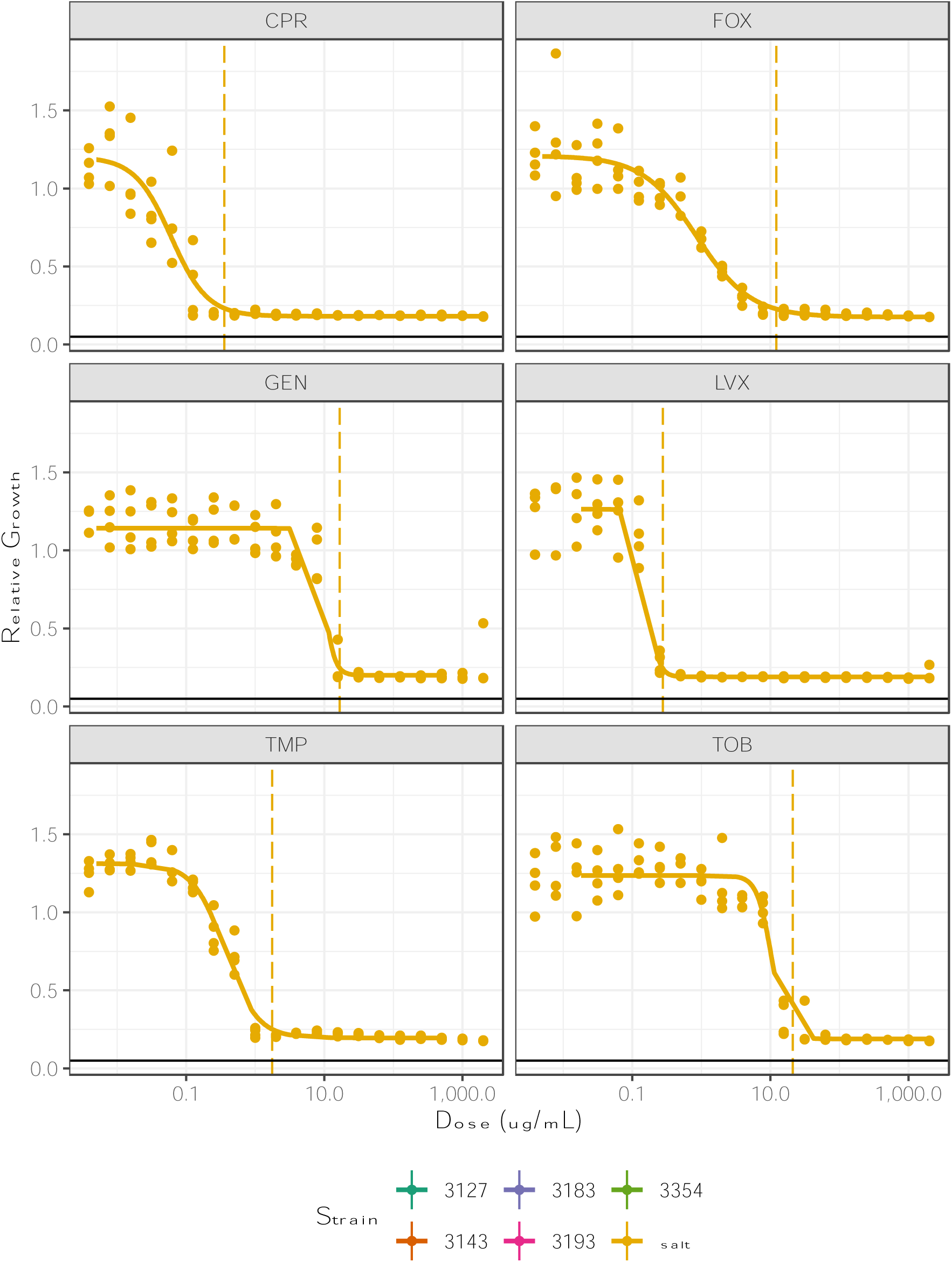
Dose response curves for multiple *E. Coli* strains grown in saltwater and in freshwater LB media for a subset of drugs tested. Colors differentiate strains and vertical lines indicate MIC 95 values. While there are differences between all the strains, the effect of saltwater cannot be determined in detectable ways, indicating that the MIC values are not impacted in meaningful ways.

**A6.**
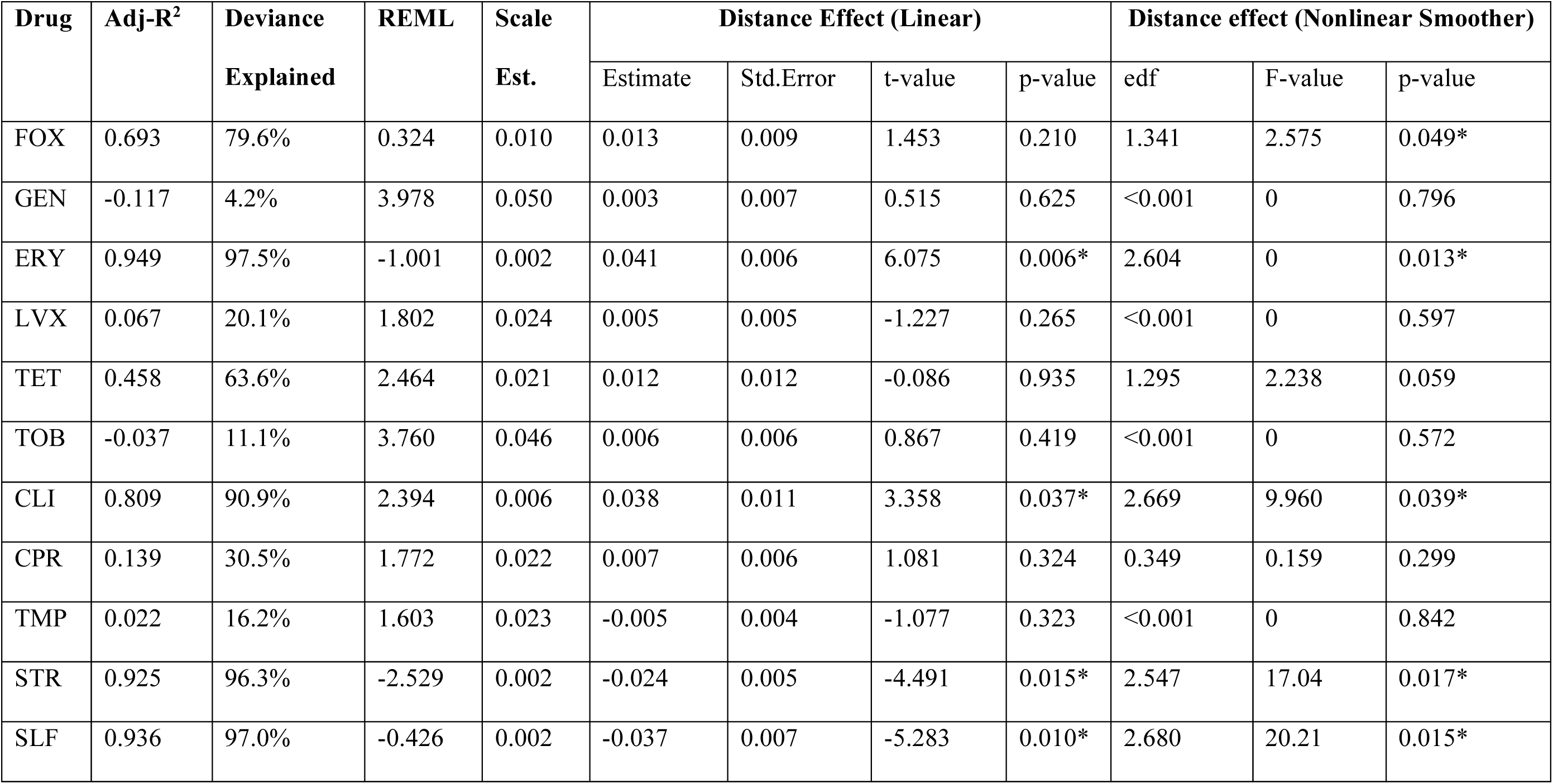
GAM output of relative MIC vs Distance (*S4*). For each drug, n = 32 and the smoother parameter reference degrees of freedom is 3. Asterisks indicate a significant effect (p-value <0.05).

| Drug | Adj-R<sup>2</sup> | Deviance<br>Explained | REML | Scale<br>Est. | Distance Effect (Linear) |  |  |  | Distance effect (Nonlinear Smoother) |  |  |
| --- | --- | --- | --- | --- | --- | --- | --- | --- | --- | --- | --- |
|  |  |  |  |  | Estimate | Std.Error | t-value | p-value | edf | F-value | p-value |
| FOX | 0.693 | 79.6% | 0.324 | 0.010 | 0.013 | 0.009 | 1.453 | 0.210 | 1.341 | 2.575 | 0.049* |
| GEN | -0.117 | 4.2% | 3.978 | 0.050 | 0.003 | 0.007 | 0.515 | 0.625 | <0.001 | 0 | 0.796 |
| ERY | 0.949 | 97.5% | -1.001 | 0.002 | 0.041 | 0.006 | 6.075 | 0.006* | 2.604 | 0 | 0.013* |
| LVX | 0.067 | 20.1% | 1.802 | 0.024 | 0.005 | 0.005 | -1.227 | 0.265 | <0.001 | 0 | 0.597 |
| TET | 0.458 | 63.6% | 2.464 | 0.021 | 0.012 | 0.012 | -0.086 | 0.935 | 1.295 | 2.238 | 0.059 |
| TOB | -0.037 | 11.1% | 3.760 | 0.046 | 0.006 | 0.006 | 0.867 | 0.419 | <0.001 | 0 | 0.572 |
| CLI | 0.809 | 90.9% | 2.394 | 0.006 | 0.038 | 0.011 | 3.358 | 0.037* | 2.669 | 9.960 | 0.039* |
| CPR | 0.139 | 30.5% | 1.772 | 0.022 | 0.007 | 0.006 | 1.081 | 0.324 | 0.349 | 0.159 | 0.299 |
| TMP | 0.022 | 16.2% | 1.603 | 0.023 | -0.005 | 0.004 | -1.077 | 0.323 | <0.001 | 0 | 0.842 |
| STR | 0.925 | 96.3% | -2.529 | 0.002 | -0.024 | 0.005 | -4.491 | 0.015* | 2.547 | 17.04 | 0.017* |
| SLF | 0.936 | 97.0% | -0.426 | 0.002 | -0.037 | 0.007 | -5.283 | 0.010* | 2.680 | 20.21 | 0.015* |

**A7.** GAM output of MIC vs Distance (*Fig.2B*). For each drug, n = 32 and the smoother reference degrees of freedom is 3. Asterisks indicate a significant effect (p-value <0.05).

| Drug | Adj-R<sup>2</sup> | Deviance<br>Explained | REML | Scale<br>Est. | Distance Effect (Linear) |  |  |  | Distance effect (Nonlinear Smoother) |  |  |
| --- | --- | --- | --- | --- | --- | --- | --- | --- | --- | --- | --- |
|  |  |  |  |  | Estimate | Std.Error | t-value | p-value | edf | F-value | p-value |
| FOX | 0.701 | 73.3% | 8.349 | 0.056 | 0.062 | 0.015 | 4.014 | <0.001* | 2.347 | 12.78 | <0.001* |
| GEN | 0.555 | 60.9% | 15.199 | 0.080 | 0.078 | 0.019 | 3.998 | <0.001* | 2.735 | 8.97 | <0.001* |
| ERY | 0.761 | 79% | 15.374 | 0.081 | 0.112 | 0.020 | 5.74 | <0.001* | 2.724 | 12.56 | <0.001* |
| LVX | 0.277 | 35.4% | 27.073 | 0.197 | 0.084 | 0.029 | 2.925 | 0.007* | 2.315 | 4.231 | 0.005* |
| TET | 0.876 | 89.1% | 26.341 | 0.162 | 0.184 | 0.028 | 6.603 | <0.001* | 2.813 | 52.86 | <0.001* |
| TOB | 0.76 | 79% | 0.394 | 0.027 | 0.087 | 0.012 | 7.531 | <0.001* | 2.884 | 23.97 | <0.001* |
| CLI | 0.348 | 41.3% | 29.862 | 0.246 | 0.037 | 0.030 | 1.213 | 0.235 | 2.08 | 5.841 | <0.001* |
| CPR | 0.237 | 29.6% | 16.415 | 0.109 | 0.019 | 0.016 | 1.239 | 0.225 | 1.41 | 2.712 | 0.007* |
| TMP | 0.291 | 35% | 48.656 | 0.916 | 0.020 | 0.049 | 0.416 | 0.680 | 1.571 | 4.494 | 0.001* |
| STR | -0.033 | 0.00% | 29.412 | 0.282 | <0.001 | <0.001 | -0.009 | 0.993 | <0.001 | 0 | 0.427 |
| SLF | 0.741 | 76.8% | 31.308 | 0.265 | -0.020 | 0.032 | -0.614 | 0.544 | 2.212 | 20.68 | <0.001* |

**A8.** GAM output of MIC vs rescaled Shannon Diversity Index (*Fig. 6*). For each drug, n = 32 and the smoother reference degrees of freedom is 3. Asterisks indicate a significant effect (p-value <0.05).

| Drug | Adj-R<sup>2</sup> | Deviance<br>Explained | REML | Scale<br>Est. | Distance Effect (Linear) |  |  |  | Distance effect (Nonlinear Smoother) |  |  |
| --- | --- | --- | --- | --- | --- | --- | --- | --- | --- | --- | --- |
|  |  |  |  |  | Estimate | Std.Error | t-value | p-value | edf | F-value | p-value |
| FOX | 0.709 | 74% | 4.478 | 0.055 | 1.006 | 0.404 | 2.493 | 0.019* | 2.297 | 15.91 | <0.001* |
| GEN | 0.696 | 73.1% | 5.217 | 0.055 | 2.472 | 0.425 | 5.816 | <0.001* | 2.547 | 9.293 | <0.001* |
| ERY | 0.309 | 38.2% | 26.285 | 0.234 | 2.745 | 0.834 | 3.293 | 0.003* | 2.284 | 4.499 | 0.004* |
| LVX | 0.330 | 35.2% | 19.536 | 0.182 | 0.786 | 0.195 | 4.034 | <0.001* | <0.001 | 0 | 0.919 |
| TET | 0.901 | 91.2% | 18.577 | 0.129 | 4.013 | 0.666 | 6.025 | <0.001* | 2.651 | 43.48 | <0.001* |
| TOB | 0.728 | 76% | -3.099 | 0.031 | 1.653 | 0.323 | 5.119 | <0.001* | 2.6 | 9.352 | <0.001* |
| CLI | 0.293 | 31.6% | 25.258 | 0.267 | 0.878 | 0.236 | 3.719 | <0.001* | <0.001 | 0 | 0.552 |
| CPR | 0.290 | 34.2% | 11.866 | 0.101 | 0.174 | 0.367 | 0.473 | 0.640 | 1.29 | 2.261 | 0.012* |
| TMP | 0.164 | 21.7% | 47.024 | 1.081 | 0.637 | 0.977 | 0.652 | 0.520 | 0.984 | 1.108 | 0.052 |
| STR | 0.008 | 4.01% | 25.46 | 0.270 | 0.266 | 0.238 | 1.119 | 0.272 | <0.001 | 0 | 0.344 |
| SLF | 0.627 | 66.3% | 33.105 | 0.383 | -1.568 | 0.999 | -1.569 | 0.128 | 2.045 | 14.58 | <0.001* |

**A9.** Linear model output of MIC vs qPCR for sulfamonomethoxine (SLF) and tetracycline (TET)

| Drug | Adj R<sup>2</sup> | Estimate | Std. Error | t value | p value |
| --- | --- | --- | --- | --- | --- |
| SLF | 0.510 | 46.25 | 8.28 | 5.586 | <0.001 |
| TET | 0.340 | -653.83 | 190.35 | -3.435 | 0.003 |

**A10.** Linear model output of Tmax for bacterial communities at site.

A9 - Linear model output of MIC vs qPCR for sulfamonomethoxine (SLF) and tetracycline (TET)
| Coefficient | Estimate | Std. Error | t-value | p-value |
| --- | --- | --- | --- | --- |
| Intercept | 53.331 | 0.828 | 64.388 | <0.001 |
| Site 2 | 0.807 | 1.265 | 0.638 | 0.530 |
| Site 3 | 1.959 | 1.171 | 1.672 | 0.108 |
| Site 4 | -0.952 | 1.171 | -0.813 | 0.425 |
| Site 5 | -1.479 | 1.171 | -1.263 | 0.219 |
| Site 6 | -2.208 | 1.171 | -1.885 | 0.072 |
| Site 7 | 0.618 | 1.171 | 0.527 | 0.603 |
| Site 8 | -2.413 | 1.171 | -2.060 | 0.051 |

**A11.** Linear model output of Topt for bacterial communities at each site.

A10 - Linear model output of Tmax for bacterial communities at site
| Coefficient | Estimate | Std. Error | t-value | p-value |
| --- | --- | --- | --- | --- |
| Intercept | 36.651 | 1.559 | 23.503 | <0.001 |
| Site 2 | -4.172 | 2.382 | -1.752 | 0.093 |
| Site 3 | -3.397 | 2.205 | -1.540 | 0.137 |
| Site 4 | -4.314 | 2.205 | -1.956 | 0.063 |
| Site 5 | -4.022 | 2.205 | -1.824 | 0.081 |
| Site 6 | -3.550 | 2.205 | -1.609 | 0.121 |
| Site 7 | -5.158 | 2.205 | -2.339 | 0.028 |
| Site 8 | -3.845 | 2.205 | -1.743 | 0.095 |

**A12.** Linear model output of Tmin for bacterial communities at each site Primer sequences.

*A11* - Linear model output of T<sub>opt</sub> for bacterial communities at each site
| Coefficient | Estimate | Std. Error | t-value | p-value |
| --- | --- | --- | --- | --- |
| Intercept | 9.797 | 0.708 | 13.829 | <0.001 |
| Site 2 | -0.730 | 1.082 | -0.675 | 0.507 |
| Site 3 | -2.690 | 1.002 | -2.685 | 0.013 |
| Site 4 | -1.193 | 1.002 | -1.191 | 0.246 |
| Site 5 | -1.308 | 1.002 | -1.306 | 0.205 |
| Site 6 | -0.821 | 1.002 | -0.819 | 0.421 |
| Site 7 | -1.444 | 1.002 | -1.442 | 0.163 |
| Site 8 | -1.919 | 1.002 | -1.915 | 0.068 |

**A13.**
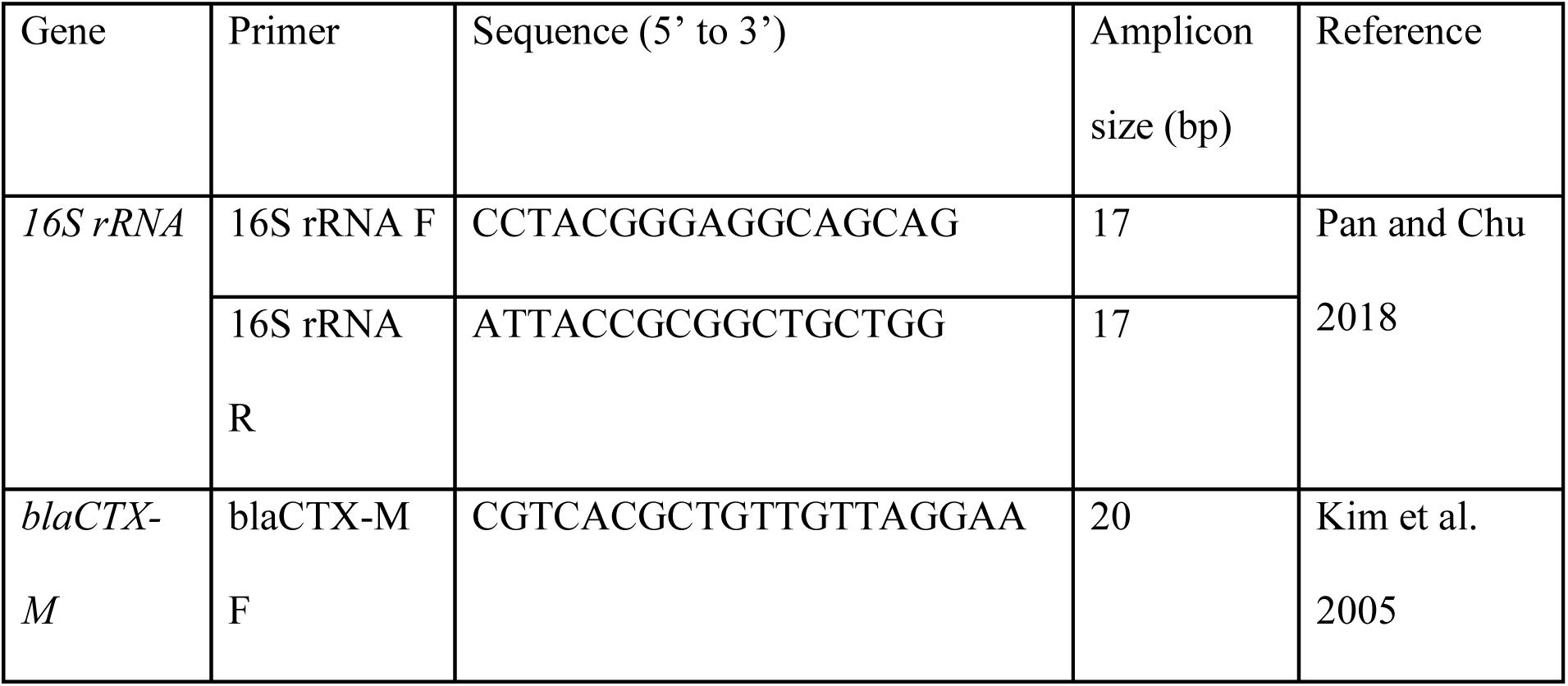
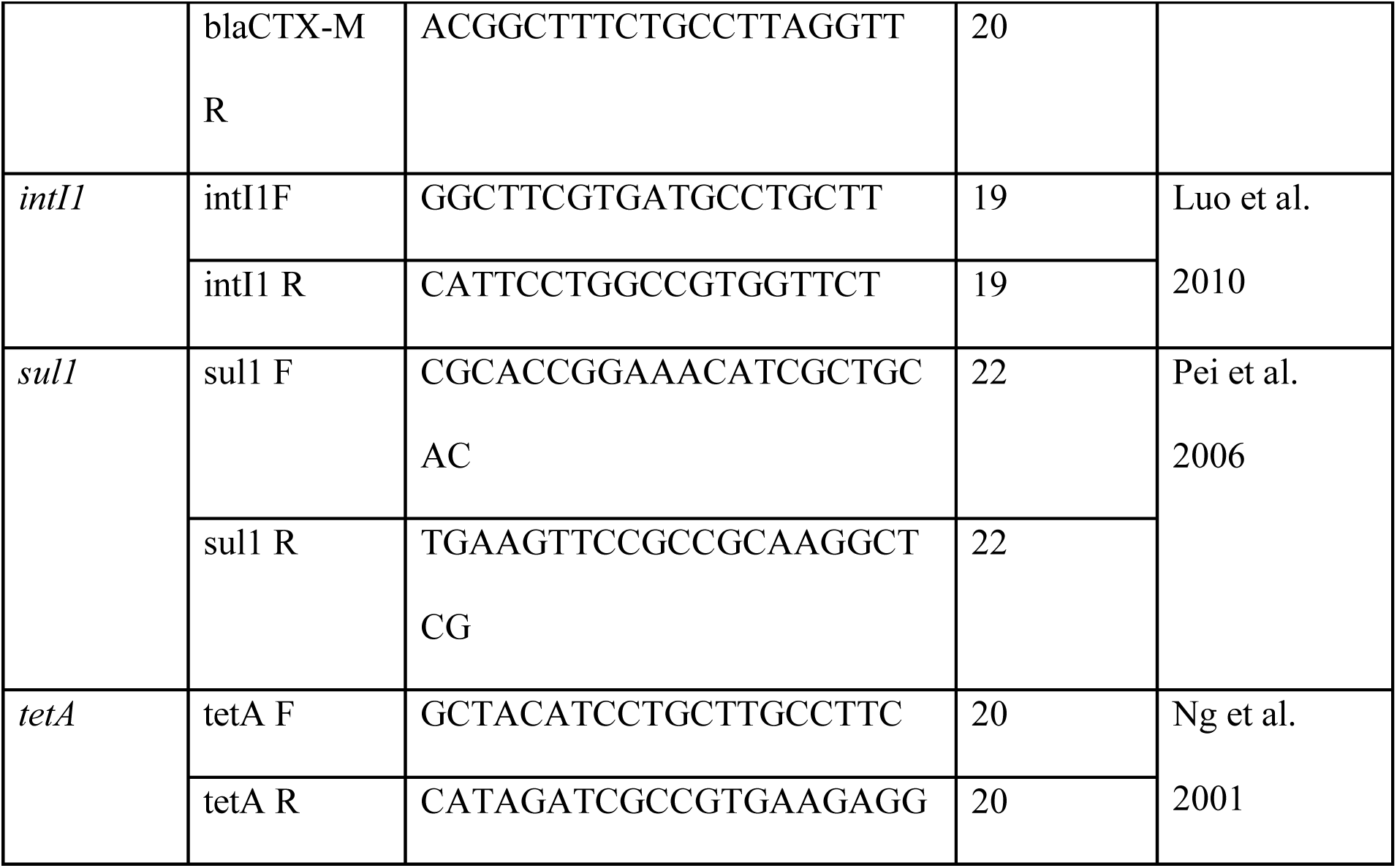
Table for all primers and sequences used for qPCR analysis. All primers were resuspended into stock solutions of a concentration of 100 uM, per manufacturer’s instructions, then diluted to a working concentration of 4 uM when loaded into well plates.

**A14.**
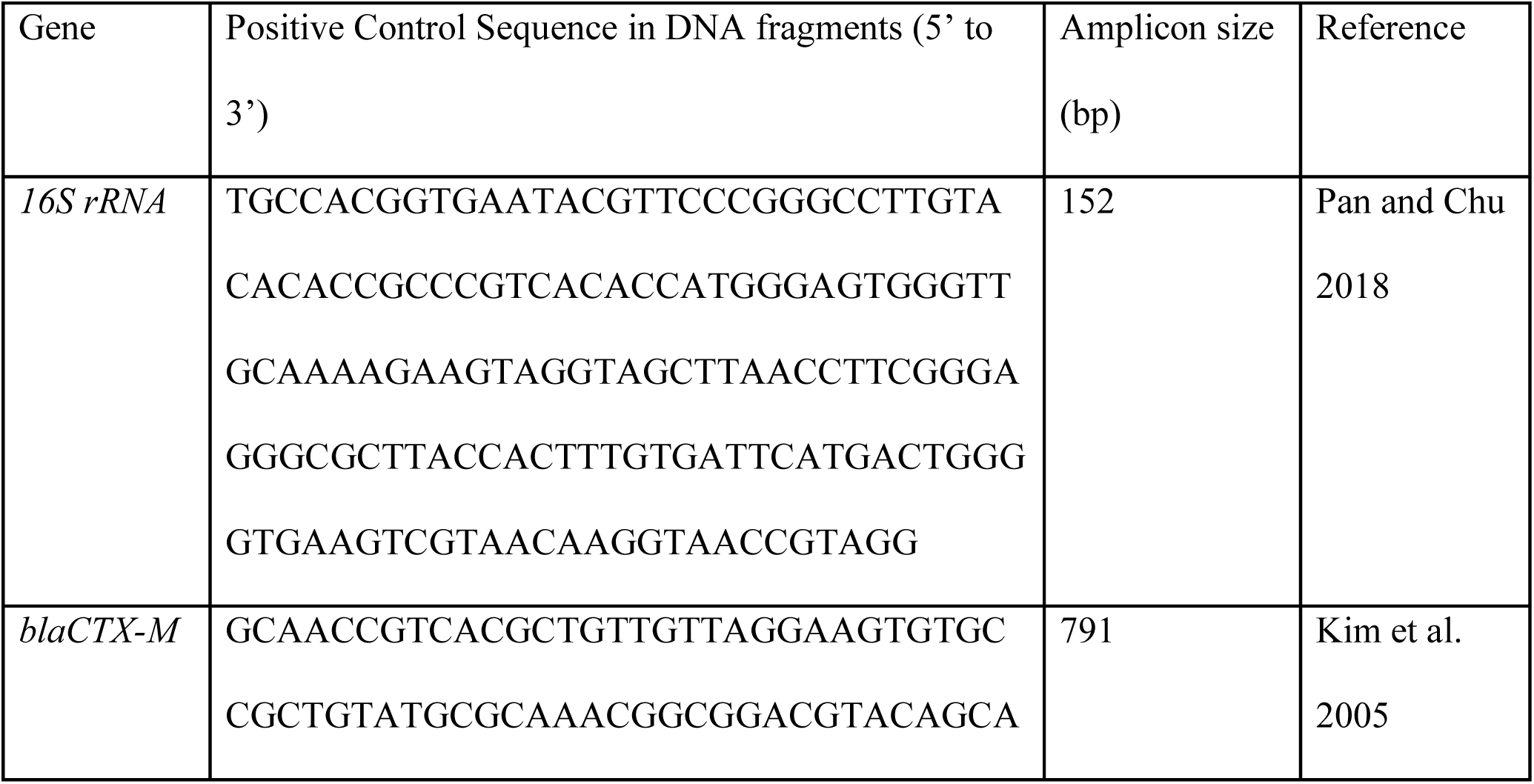
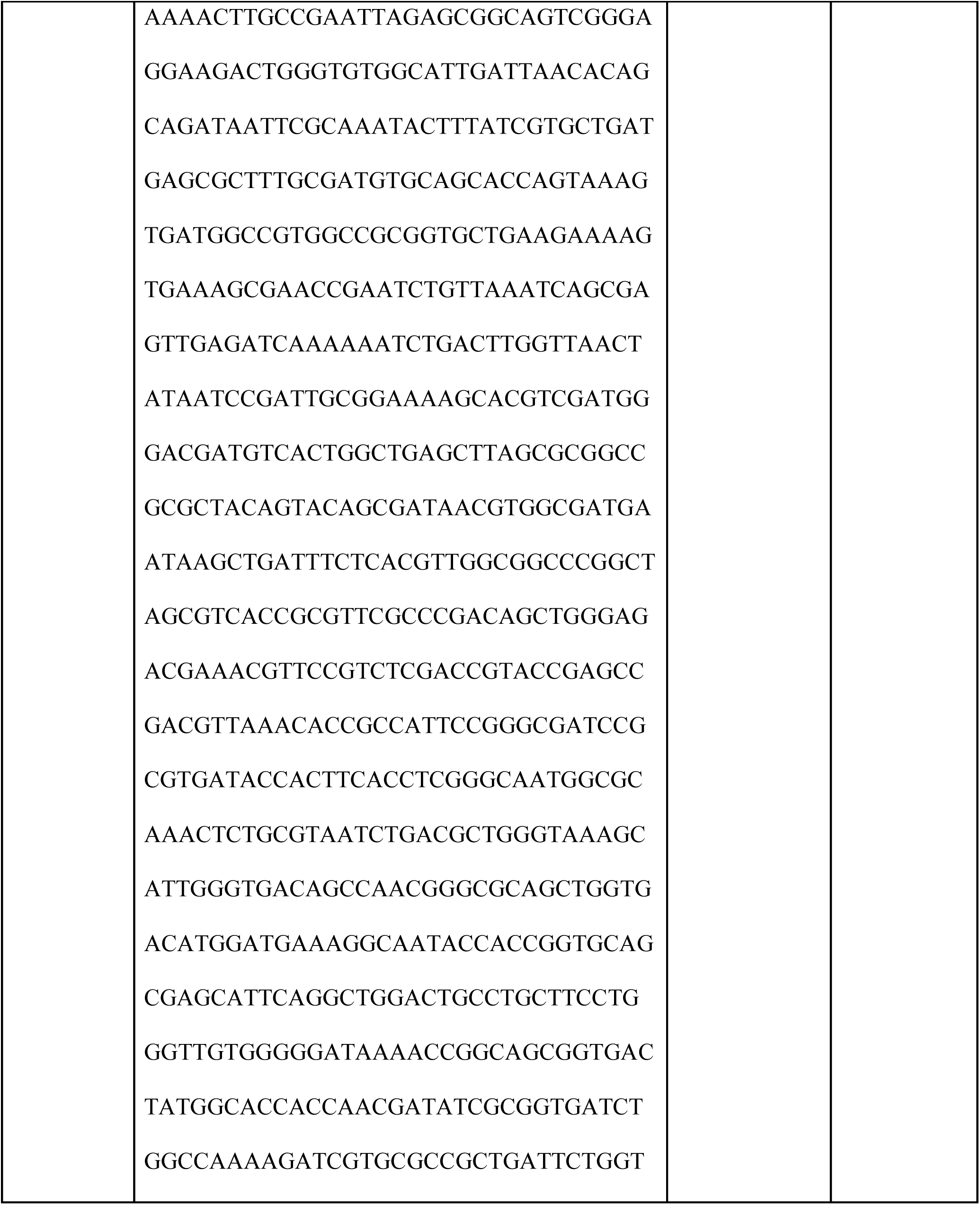
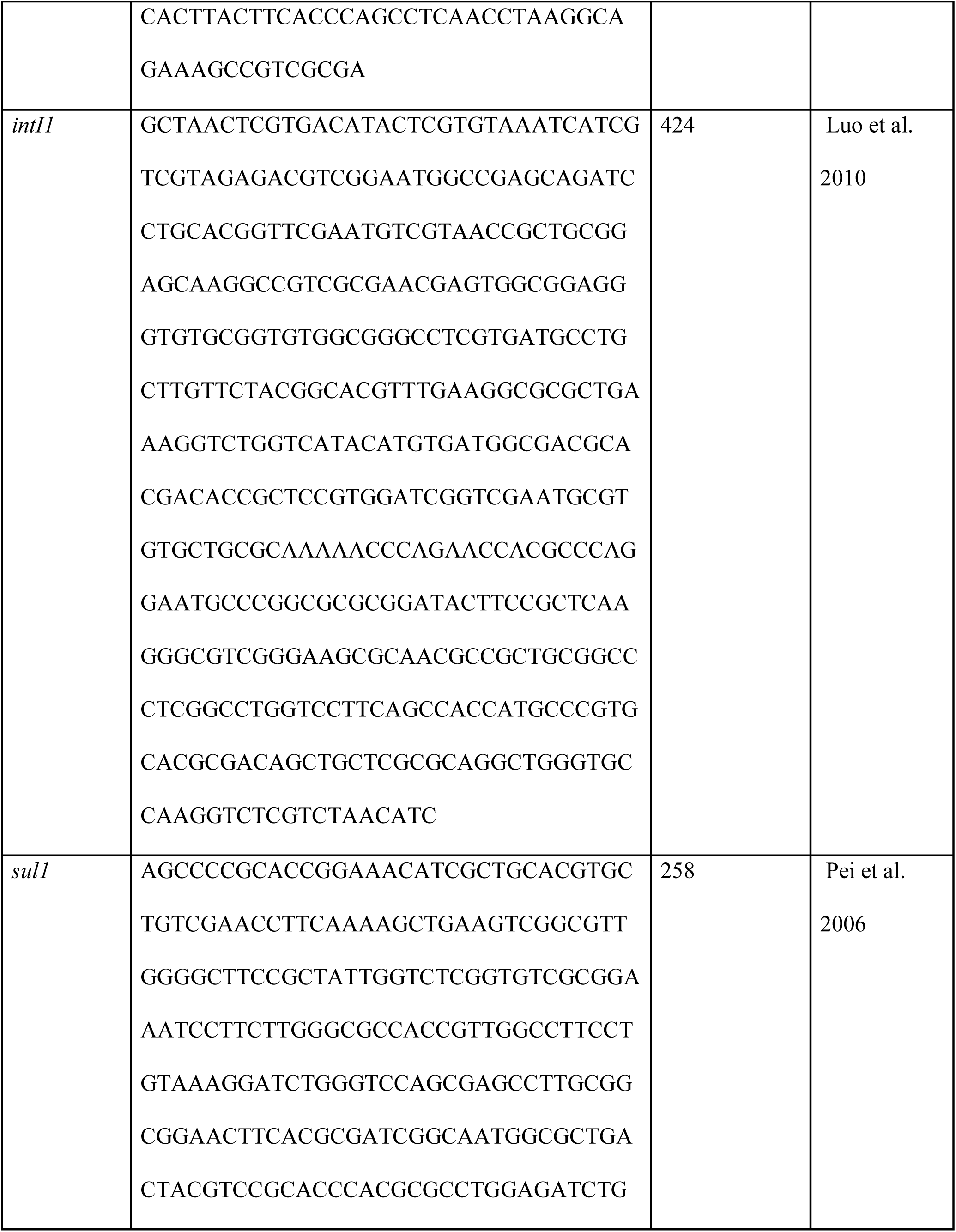
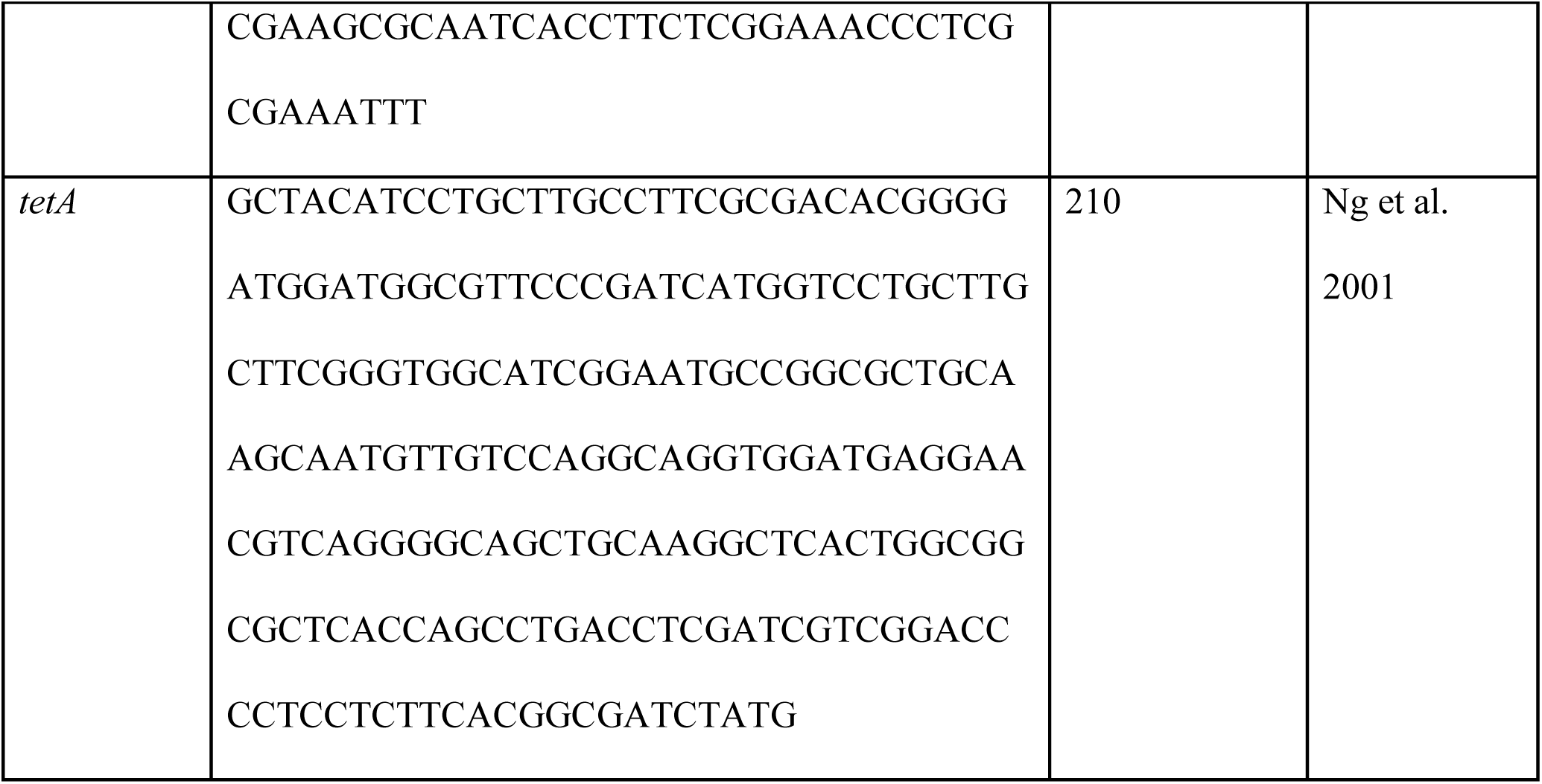
Table for positive control sequences for genes being amplified to compare gene quantities in bacterial communities at each site.

**A15.** Volumes of each reagent loaded into each well in 96-well plates used for qPCR.

| Reagent | Volume (uL) |
| --- | --- |
| PowerUp SYBR Green MasterMix | 12.5 |
| Forward primer | 1 |
| Reverse primer | 1 |
| Molecular-grade water | 2.5 |
| Template DNA | 3 |

## References

1. Aguiló-Ferretjans MM, Bosch R, Martín-Cardona C, Lalucat J, Nogales B. 2008. Phylogenetic analysis of the composition of bacterial communities in human-exploited coastal environments from Mallorca Island (Spain). Systematic and Applied Microbiology 31:231–240.

2. Allen HK, Donato J, Wang HH, Cloud-Hansen KA, Davies J, Handelsman J. 2010. Call of the wild: antibiotic resistance genes in natural environments. Nat Rev Microbiol 8:251–259.

3. Almakki A, Jumas-Bilak E, Marchandin H, Licznar-Fajardo P. 2019. Antibiotic resistance in urban runoff. Science of The Total Environment 667:64–76.

4. Amarasiri M, Sano D, Suzuki S. 2020. Understanding human health risks caused by antibiotic resistant bacteria (ARB) and antibiotic resistance genes (ARG) in water environments: Current knowledge and questions to be answered. Critical Reviews in Environmental Science and Technology 50:2016–2059.

5. Aminov RI. 2009. The role of antibiotics and antibiotic resistance in nature. Environmental Microbiology 11:2970–2988.

6. Aminov RI. 2011. Horizontal Gene Exchange in Environmental Microbiota. Front Microbiol 2:158.

7. Antunes P, Machado J, Sousa JC, Peixe L. 2005. Dissemination of Sulfonamide Resistance Genes (*sul1*, *sul2*, and *sul3*) in Portuguese *Salmonella enterica* Strains and Relation with Integrons. Antimicrob Agents Chemother 49:836–839.

8. Baya AM, Brayton PR, Brown VL, Grimes DJ, Russek-Cohen E, Colwell RR. 1986. Coincident plasmids and antimicrobial resistance in marine bacteria isolated from polluted and unpolluted Atlantic Ocean samples. Appl Environ Microbiol 51:1285–1292.

9. Blondeau JM. 2004. Fluoroquinolones: mechanism of action, classification, and development of resistance. Survey of Ophthalmology 49:S73–S78.

10. Brown C, Corcoran E, Herkenrath P. 2006. Marine and coastal ecosystems and human well-being: A synthesis report based on the findings of the Millennium Ecosystem Assessment.

11. Bullivant A, Lozano-Huntelman N, Tabibian K, Leung V, Armstrong D, Dudley H, Savage VM, Rodríguez-Verdugo A, Yeh PJ. 2024. Evolution Under Thermal Stress Affects *Escherichia coli* ’s Resistance to Antibiotics 10.1101/2024.02.27.582334.

12. Cáceres MD, Legendre P. 2009. Associations between species and groups of sites: indices and statistical inference. Ecology 90:3566–3574.

13. Chen Q-L, An X-L, Zheng B-X, Gillings M, Peñuelas J, Cui L, Su J-Q, Zhu Y-G. 2019. Loss of soil microbial diversity exacerbates spread of antibiotic resistance. Soil Ecol Lett 1:3–13.

14. Chow LKM, Ghaly TM, Gillings MR. 2021. A survey of sub-inhibitory concentrations of antibiotics in the environment. Journal of Environmental Sciences 99:21–27.

15. Costanzo SD, Murby J, Bates J. 2005. Ecosystem response to antibiotics entering the aquatic environment. Marine Pollution Bulletin 51:218–223.

16. Cottrell MT, Kirchman DL. 2000. Community Composition of Marine Bacterioplankton Determined by 16S rRNA Gene Clone Libraries and Fluorescence In Situ Hybridization. Applied and Environmental Microbiology 66:5116–5122.

17. Cram JA, Chow C-ET, Sachdeva R, Needham DM, Parada AE, Steele JA, Fuhrman JA. 2015. Seasonal and interannual variability of the marine bacterioplankton community throughout the water column over ten years. 3. ISME J 9:563–580.

18. Cruz-Loya M, Kang TM, Lozano NA, Watanabe R, Tekin E, Damoiseaux R, Savage VM, Yeh PJ. 2019. Stressor interaction networks suggest antibiotic resistance co-opted from stress responses to temperature. 1. ISME J 13:12–23.

19. Cuadrat RRC, Sorokina M, Andrade BG, Goris T, Dávila AMR. 2020. Global ocean resistome revealed: Exploring antibiotic resistance gene abundance and distribution in TARA Oceans samples. GigaScience 9:giaa046.

20. D’Costa VM, McGrann KM, Hughes DW, Wright GD. 2006. Sampling the Antibiotic Resistome. Science 311:374–377.

21. D’Costa VM, King CE, Kalan L, Morar M, Sung WWL, Schwarz C, Froese D, Zazula G, Calmels F, Debruyne R, Golding GB, Poinar HN, Wright GD. 2011. Antibiotic resistance is ancient. Nature 477:457–461.

22. Echeverria-Palencia CM, Thulsiraj V, Tran N, Ericksen CA, Melendez I, Sanchez MG, Walpert D, Yuan T, Ficara E, Senthilkumar N, Sun F, Li R, Hernandez-Cira M, Gamboa D, Haro H, Paulson SE, Zhu Y, Jay JA. 2017. Disparate Antibiotic Resistance Gene Quantities Revealed across 4 Major Cities in California: A Survey in Drinking Water, Air, and Soil at 24 Public Parks. ACS Omega 2:2255–2263.

23. Fernández-Villa D, Aguilar MR, Rojo L. 2019. Folic Acid Antagonists: Antimicrobial and Immunomodulating Mechanisms and Applications. 20. International Journal of Molecular Sciences 20:4996.

24. Finley RL, Collignon P, Larsson DGJ, McEwen SA, Li X-Z, Gaze WH, Reid-Smith R, Timinouni M, Graham DW, Topp E. 2013. The Scourge of Antibiotic Resistance: The Important Role of the Environment. Clinical Infectious Diseases 57:704–710.

25. Forge A, Schacht J. 2000. Aminoglycoside Antibiotics. Audiol Neurotol 5:3–22.

26. Fortunato CS, Herfort L, Zuber P, Baptista AM, Crump BC. 2012. Spatial variability overwhelms seasonal patterns in bacterioplankton communities across a river to ocean gradient. 3. ISME J 6:554–563.

27. Fresia P, Antelo V, Salazar C, Giménez M, D’Alessandro B, Afshinnekoo E, Mason C, Gonnet GH, Iraola G. 2019. Urban metagenomics uncover antibiotic resistance reservoirs in coastal beach and sewage waters. Microbiome 7:35.

28. Frieri M, Kumar K, Boutin A. 2017. Antibiotic resistance. Journal of Infection and Public Health 10:369–378.

29. Fuhrman JA, Cram JA, Needham DM. 2015. Marine microbial community dynamics and their ecological interpretation. Nat Rev Microbiol 13:133–146.

30. Fuhrman JA, Hewson I, Schwalbach MS, Steele JA, Brown MV, Naeem S. 2006. Annually reoccurring bacterial communities are predictable from ocean conditions. Proceedings of the National Academy of Sciences 103:13104–13109.

31. Gillings MR, Gaze WH, Pruden A, Smalla K, Tiedje JM, Zhu Y-G. 2015. Using the class 1 integron-integrase gene as a proxy for anthropogenic pollution. 6. ISME J 9:1269–1279.

32. Given S, Pendleton LH, Boehm AB. 2006. Regional Public Health Cost Estimates of Contaminated Coastal Waters: A Case Study of Gastroenteritis at Southern California Beaches. Environ Sci Technol 40:4851–4858.

33. Goldberg N. 2023. Long Beach closes beaches ahead of a warm weekend after L.A. River sewage spill. Los Angeles Times.

34. Gross S, Müller A, Seinige D, Wohlsein P, Oliveira M, Steinhagen D, Kehrenberg C, Siebert U. 2022. Occurrence of Antimicrobial-Resistant Escherichia coli in Marine Mammals of the North and Baltic Seas: Sentinels for Human Health. Antibiotics 11:1248.

35. Grossman TH. 2016. Tetracycline Antibiotics and Resistance. Cold Spring Harb Perspect Med 6:a025387.

36. Gumprecht B. 1997. 51 Miles of Concrete: The Exploitation and Transformation of the Los Angeles River. Southern California Quarterly 79:431–486.

37. Halling-Sørensen B, Nors Nielsen S, Lanzky PF, Ingerslev F, Holten Lützhøft HC, Jørgensen SE. 1998. Occurrence, fate and effects of pharmaceutical substances in the environment-A review. Chemosphere 36:357–393.

38. Hatosy SM, Martiny AC. 2015. The Ocean as a Global Reservoir of Antibiotic Resistance Genes. Applied and Environmental Microbiology 81:7593–7599.

39. Hatosy SM, Martiny JBH, Sachdeva R, Steele J, Fuhrman JA, Martiny AC. 2013. Beta diversity of marine bacteria depends on temporal scale. Ecology 94:1898–1904.

40. Hijmans R. 2022. geosphere: Spherical Trigonometry (R package version 1.5-18).

41. Hossain A, Habibullah-Al-Mamun Md, Nagano I, Masunaga S, Kitazawa D, Matsuda H. 2022. Antibiotics, antibiotic-resistant bacteria, and resistance genes in aquaculture: risks, current concern, and future thinking. Environ Sci Pollut Res 29:11054–11075.

42. Hutchings MI, Truman AW, Wilkinson B. 2019. Antibiotics: past, present and future. Current Opinion in Microbiology 51:72–80.

43. Iwu CD, Korsten L, Okoh AI. 2020. The incidence of antibiotic resistance within and beyond the agricultural ecosystem: A concern for public health. MicrobiologyOpen 9:e1035.

44. Ju F, Beck K, Yin X, Maccagnan A, McArdell CS, Singer HP, Johnson DR, Zhang T, Bürgmann H. 2019. Wastewater treatment plant resistomes are shaped by bacterial composition, genetic exchange, and upregulated expression in the effluent microbiomes. The ISME Journal 13:346– 360.

45. Kamruzzaman M, Iredell J. 2019. A ParDE-family toxin antitoxin system in major resistance plasmids of Enterobacteriaceae confers antibiotic and heat tolerance. Sci Rep 9:9872.

46. Knapp CW, Dolfing J, Ehlert PAI, Graham DW. 2010. Evidence of Increasing Antibiotic Resistance Gene Abundances in Archived Soils since 1940. Environ Sci Technol 44:580–587.

47. Kowalska-Krochmal B, Dudek-Wicher R. 2021. The Minimum Inhibitory Concentration of Antibiotics: Methods, Interpretation, Clinical Relevance. Pathogens 10:165.

48. Kraemer SA, Ramachandran A, Perron GG. 2019. Antibiotic Pollution in the Environment: From Microbial Ecology to Public Policy. Microorganisms 7:180.

49. Kremer C. 2023. growthTools: Tools for analyzing time series of microbial abundances to estimate growth rates (R package version 0.1.2).

50. Langenheder S, Lindström ES. 2019. Factors influencing aquatic and terrestrial bacterial community assembly. Environ Microbiol Rep 11:306–315.

51. Larsson DGJ, Flach C-F. 2022. Antibiotic resistance in the environment. 5. Nat Rev Microbiol 20:257–269.

52. Lartigue M-F, Poirel L, Nordmann P. 2004. Diversity of genetic environment of *bla* CTX-M genes. FEMS Microbiology Letters 234:201–207.

53. Leonard AFC, Zhang L, Balfour AJ, Garside R, Hawkey PM, Murray AK, Ukoumunne OC, Gaze WH. 2018. Exposure to and colonisation by antibiotic-resistant E. coli in UK coastal water users: Environmental surveillance, exposure assessment, and epidemiological study (Beach Bum Survey). Environment International 114:326–333.

54. Li R, Jay JA, Stenstrom MK. 2019. Fate of antibiotic resistance genes and antibiotic-resistant bacteria in water resource recovery facilities. Water Environ Res 91:5–20.

55. Liao J, Chen Y. 2018. Removal of intl1 and associated antibiotics resistant genes in water, sewage sludge and livestock manure treatments. Rev Environ Sci Biotechnol 17:471–500.

56. Livermore DM. 2009. Has the era of untreatable infections arrived? Journal of Antimicrobial Chemotherapy 64:i29–i36.

57. Logares R, Bråte J, Bertilsson S, Clasen JL, Shalchian-Tabrizi K, Rengefors K. 2009. Infrequent marine–freshwater transitions in the microbial world. Trends in Microbiology 17:414–422.

58. Manaia CM. 2017. Assessing the Risk of Antibiotic Resistance Transmission from the Environment to Humans: Non-Direct Proportionality between Abundance and Risk. Trends in Microbiology 25:173–181.

59. McDaniel LD, Young EC, Ritchie KB, Paul JH. 2012. Environmental Factors Influencing Gene Transfer Agent (GTA) Mediated Transduction in the Subtropical Ocean. PLOS ONE 7:e43506.

60. McDaniel LD, Young E, Delaney J, Ruhnau F, Ritchie KB, Paul JH. 2010. High Frequency of Horizontal Gene Transfer in the Oceans. Science 330:50–50.

61. Mobilian C, Wisnoski NI, Lennon JT, Alber M, Widney S, Craft CB. 2020. Differential effects of press vs. pulse seawater intrusion on microbial communities of a tidal freshwater marsh. Limnol Oceanogr Letters 8:154–161.

62. Møller TSB, Overgaard M, Nielsen SS, Bortolaia V, Sommer MOA, Guardabassi L, Olsen JE. 2016. Relation between tetR and tetA expression in tetracycline resistant Escherichia coli. BMC Microbiol 16:39.

63. Mulder CPH, Uliassi DD, Doak DF. 2001. Physical stress and diversity-productivity relationships: The role of positive interactions. Proc Natl Acad Sci USA 98:6704–6708.

64. Munita JM, Arias CA. 2016. Mechanisms of Antibiotic Resistance. Microbiol Spectr 4:4.2.15.

65. Neu HC. 1976. Tobramycin: An Overview. Journal of Infectious Diseases 134:S3–S19.

66. Nielsen MC, Wang N, Jiang SC. 2021. Acquisition of antibiotic resistance genes on human skin after swimming in the ocean. Environmental Research 197:110978.

67. Niu Z-G, Zhang K, Zhang Y. 2016. Occurrence and distribution of antibiotic resistance genes in the coastal area of the Bohai Bay, China. Marine Pollution Bulletin 107:245–250.

68. Nogales B, Aguiló-Ferretjans MM, Martín-Cardona C, Lalucat J, Bosch R. 2007. Bacterial diversity, composition and dynamics in and around recreational coastal areas. Environmental Microbiology 9:1913–1929.

69. Nogales B, Lanfranconi MP, Piña-Villalonga JM, Bosch R. 2011. Anthropogenic perturbations in marine microbial communities. FEMS Microbiology Reviews 35:275–298.

70. Norberg J. 2004. Biodiversity and ecosystem functioning: A complex adaptive systems approach. Limnology & Oceanography 49:1269–1277.

71. Oates JA, Wood AJJ, Donowitz GR, Mandell GL. 1988. Beta-Lactam Antibiotics. N Engl J Med 318:419–426.

72. Oksanen J, Simpson G, Blanchet G, Kindt R, Legendre P, Minchin P, O’Hara RB, Solymos P, Stevens H, Szoecs E, Wagner H, Barbour M, Bedward M, Bolker B, Borcard D, Carvalho G, Chirico M, De Caceres M, Durand S, Evangelista H, FitzJohn R, Friendly M, Furneaux B, Hannigan G, Hill M, Lahti L, McGlinn D, Ouellette M-H, Ribeiro Cunha E, Smith T, Stier A, Ter Braak C, Weedon J. 2024. vegan: Community Ecology Package.

73. Paver SF, Muratore D, Newton RJ, Coleman ML. 2018. Reevaluating the Salty Divide: Phylogenetic Specificity of Transitions between Marine and Freshwater Systems. mSystems 3:e00232–18.

74. Pruden A, Larsson DGJ, Amézquita A, Collignon P, Brandt KK, Graham DW, Lazorchak JM, Suzuki S, Silley P, Snape JR, Topp E, Zhang T, Zhu Y-G. 2013. Management Options for Reducing the Release of Antibiotics and Antibiotic Resistance Genes to the Environment. Environmental Health Perspectives 121:878–885.

75. R Core Team. 2024. R: A Language and Environment for Statistical Computing. R Foundation for Statistical Computing, Austria, Vienna.

76. Ritz C, Baty F, Streibig JC, Gerhard D. 2015. Dose-Response Analysis Using R. PLOS ONE 10:e0146021.

77. Rizzo L, Manaia C, Merlin C, Schwartz T, Dagot C, Ploy MC, Michael I, Fatta-Kassinos D. 2013. Urban wastewater treatment plants as hotspots for antibiotic resistant bacteria and genes spread into the environment: A review. Science of The Total Environment 447:345–360.

78. Rodríguez-Verdugo A, Gaut BS, Tenaillon O. 2013. Evolution of Escherichia coli rifampicin resistance in an antibiotic-free environment during thermal stress. BMC Evol Biol 13:50.

79. Rodríguez-Verdugo A, Lozano-Huntelman N, Cruz-Loya M, Savage V, Yeh P. 2020. Compounding Effects of Climate Warming and Antibiotic Resistance. iScience 23:101024.

80. Rose JM, Gast RJ, Bogomolni A, Ellis JC, Lentell BJ, Touhey K, Moore M. 2009. Occurrence and patterns of antibiotic resistance in vertebrates off the Northeastern United States coast: Antibiotic resistance in coastal vertebrates. FEMS Microbiology Ecology 67:421–431.

81. Schloss PD, Westcott SL, Ryabin T, Hall JR, Hartmann M, Hollister EB, Lesniewski RA, Oakley BB, Parks DH, Robinson CJ, Sahl JW, Stres B, Thallinger GG, Van Horn DJ, Weber CF. 2009. Introducing mothur: Open-Source, Platform-Independent, Community-Supported Software for Describing and Comparing Microbial Communities. Applied and Environmental Microbiology 75:7537–7541.

82. Sepúlveda-Correa A, Daza-Giraldo LV, Polanía J, Arenas NE, Muñoz-García A, Sandoval-Figueredo AV, Vanegas J. 2021. Genes associated with antibiotic tolerance and synthesis of antimicrobial compounds in a mangrove with contrasting salinities. Marine Pollution Bulletin 171:112740.

83. Sizemore RK, Colwell RR. 1977. Plasmids Carried by Antibiotic-Resistant Marine Bacteria. Antimicrob Agents Chemother 12:373–382.

84. Sköld O. 2000. Sulfonamide resistance: mechanisms and trends. Drug Resistance Updates 3:155–160.

85. Smieja M. 1998. Current Indications for the Use of Clindamycin: A Critical Review. Canadian Journal of Infectious Diseases and Medical Microbiology 9:22–28.

86. Solo-Gabriele HM, Harwood VJ, Kay D, Fujioka RS, Sadowsky MJ, Whitman RL, Wither A, Caniça M, Carvalho Da Fonseca R, Duarte A, Edge TA, Gargaté MJ, Gunde-Cimerman N, Hagen F, McLellan SL, Nogueira Da Silva A, Novak Babič M, Prada S, Rodrigues R, Romão D, Sabino R, Samson RA, Segal E, Staley C, Taylor HD, Veríssimo C, Viegas C, Barroso H, Brandão JC. 2016. Beach sand and the potential for infectious disease transmission: observations and recommendations. J Mar Biol Ass 96:101–120.

87. Sommer MOA, Dantas G, Church GM. 2009. Functional Characterization of the Antibiotic Resistance Reservoir in the Human Microflora. Science 325:1128–1131.

88. Søraas A, Sundsfjord A, Sandven I, Brunborg C, Jenum PA. 2013. Risk Factors for Community-Acquired Urinary Tract Infections Caused by ESBL-Producing Enterobacteriaceae –A Case– Control Study in a Low Prevalence Country. PLoS One 8:e69581.

89. Spížek J, Řezanka T. 2017. Lincosamides: Chemical structure, biosynthesis, mechanism of action, resistance, and applications. Biochemical Pharmacology 133:20–28.

90. Storteboom H, Arabi M, Davis JG, Crimi B, Pruden A. 2010. Tracking Antibiotic Resistance Genes in the South Platte River Basin Using Molecular Signatures of Urban, Agricultural, And Pristine Sources. Environ Sci Technol 44:7397–7404.

91. Szekeres E, Chiriac CM, Baricz A, Szőke-Nagy T, Lung I, Soran M-L, Rudi K, Dragos N, Coman C. 2018. Investigating antibiotics, antibiotic resistance genes, and microbial contaminants in groundwater in relation to the proximity of urban areas. Environmental Pollution 236:734–744.

92. Toprak E, Veres A, Michel J-B, Chait R, Hartl DL, Kishony R. 2012. Evolutionary paths to antibiotic resistance under dynamically sustained drug selection. Nat Genet 44:101–105.

93. Vázquez-Laslop N, Mankin AS. 2018. How Macrolide Antibiotics Work. Trends in Biochemical Sciences 43:668–684.

94. Vega NM, Gore J. 2014. Collective antibiotic resistance: mechanisms and implications. Current Opinion in Microbiology 21:28–34.

95. Wickham H. 2016. ggplot2: elegant graphics for data analysisSecond edition. Springer, Switzerland.

96. Wolfand JM, Sytsma A, Hennon VL, Stein ED, Hogue TS. 2022. Dilution and Pollution: Assessing the Impacts of Water Reuse and Flow Reduction on Water Quality in the Los Angeles River Basin. ACS EST Water 2:1309–1319.

97. Wood SN. 2011. Fast Stable Restricted Maximum Likelihood and Marginal Likelihood Estimation of Semiparametric Generalized Linear Models. Journal of the Royal Statistical Society Series B: Statistical Methodology 73:3–36.

98. Xu Y, You G, Zhang M, Peng D, Jiang Z, Qi S, Yang S, Hou J. 2022. Antibiotic resistance genes alternation in soils modified with neutral and alkaline salts: interplay of salinity stress and response strategies of microbes. Science of The Total Environment 809:152246.

99. Yeh Y-C, Fuhrman JA. 2022. Contrasting diversity patterns of prokaryotes and protists over time and depth at the San-Pedro Ocean Time series. 1. ISME COMMUN 2:1–12.

100. Zhang Y-J, Hu H-W, Yan H, Wang J-T, Lam SK, Chen Q-L, Chen D, He J-Z. 2019. Salinity as a predominant factor modulating the distribution patterns of antibiotic resistance genes in ocean and river beach soils. Science of The Total Environment 668:193–203.

101. Zhu Y-G, Zhao Y, Li B, Huang C-L, Zhang S-Y, Yu S, Chen Y-S, Zhang T, Gillings MR, Su J- Q. 2017. Continental-scale pollution of estuaries with antibiotic resistance genes. Nat Microbiol 2:16270.

102. 2020. U.S. Census Bureau QuickFacts: Long Beach city, California. https://www.census.gov/quickfacts/fact/table/longbeachcitycalifornia/POP010220. Retrieved 4 June 2023.

